# Impact of functional synapse clusters on neuronal response selectivity

**DOI:** 10.1101/634220

**Authors:** Balázs B Ujfalussy, Judit K Makara

## Abstract

Clustering of functionally similar synapses in dendrites is thought to affect input-output transformation by inducing dendritic nonlinearities. However, neither the *in vivo* impact of synaptic clusters on somatic membrane potential (sVm), nor the rules of cluster formation are elucidated. We developed a computational approach to measure the effect of functional synaptic clusters on sVm response of biophysical model CA1 and L2/3 pyramidal neurons to behaviorally relevant *in vivo*-like inputs. Large-scale dendritic spatial inhomogeneities in synaptic tuning properties did influence sVm, but small synaptic clusters appearing randomly with unstructured connectivity did not. With structured connectivity, ~10-20 synapses per cluster was optimal for clustering-based tuning, but larger responses were achieved by 2-fold potentiation of the same synapses. We further show that without nonlinear amplification of the effect of random clusters, action potential-based, global plasticity rules can not generate functional clustering. Our results suggest that clusters likely form via local synaptic interactions, and have to be moderately large to impact sVm responses.

## Introduction

Processing of synaptic stimuli targeting the dendritic tree fundamentally depends on the spatio-temporal structure of the inputs: spatially distributed or asynchronous inputs are integrated linearly, whereas spatially close and synchronous inputs induce dendritic nonlinearities, such as regenerative dendritic spikes. These observations motivated the idea of functional synaptic clustering: to elicit dendritic spikes, inputs showing correlated *in vivo* activity should target nearby dendritic locations (Larkum & Nevian, 2008). Consistent with this idea, *in vivo* imaging of the activity of dendritic spines demonstrated that neighbouring synapses are co-activated more often than random (Takahashi et al., 2012; Winnubst et al., 2015; Iacaruso et al., 2017; Scholl et al., 2017; Kerlin et al., 2018) suggesting the involvement of active processes in the formation of functional clusters. However, both the relative importance of synaptic clustering compared to other factors influencing neuronal responses under *in vivo* conditions and the biophysical mechanisms leading to their formation are unknown.

In particular, the spatial scale of the synapses showing correlated activity *in vivo* has been found to be restricted to 5-10 *μ*m and involved a small number, ~2-5 dendritic spines (Takahashi et al., 2012; Iacaruso et al., 2017; Scholl et al., 2017; Kerlin et al., 2018). This is substantially less than the 10-20 inputs required to trigger dendritic spikes under in *vitro* conditions (Mel, 1993; Losonczy & Magee, 2006; Makara & Magee, 2013; Branco & Häusser, 2011) leaving the potential impact of the clusters elusive. The high background activity, characteristic for the *in vivo* states, can markedly change the integrative properties of the cell (London & Segev, 2001; Destexhe et al., 2003; Ujfalussy et al., 2018), but it is not clear how it influences the effect of functional clustering, ie., whether the facilitation of dendritic spikes (Farinella et al., 2014; Basak & Narayanan, 2018), or other effects, such as saturation (Longordo et al., 2013), shunting (Mel, 1993; Palmer et al., 2012) or increased trial-to-trial variability (Gómez-Laberge et al., 2016) are stronger.

The formation of small functional synapse clusters likely involves activity-dependent synaptic plasticity mechanism(s) (Makino & Malinow, 2011; Zhang et al., 2015). Synaptic plasticity is influenced by both global (i.e. cell-wide) and local (i.e. dendritic) processes, and how these factors interact to generate functional clustering is not known. On one hand, several lines of evidence indicate that clusters can be generated through local plasticity mechanisms. First, synaptic plasticity has been shown to be driven by local synchrony of the inputs (Winnubst et al., 2015). Second, local cooperative synaptic plasticity mechanisms have been described in dendrites acting independently of the somatic output of the neuron on the spatial scale of synaptic clustering, i.e., 5-10 *μ*m (Harvey & Svoboda, 2007; Weber et al., 2016). Finally functional clustering was not restricted to inputs showing correlated activity with the soma (Scholl et al., 2017). On the other hand, theoretical considerations argue (Hebb, 1949; Lillicrap et al., 2016) and *in vivo* experimental evidence demonstrates (Pawlak et al., 2013) that the somatic output of the neuron influences plasticity of the incoming synapses. Global plasticity may also lead to functional synaptic clustering if accidental initial clustering of co-active synapses could facilitate somatic action potentials that, back-propagating to the dendritic tree, can reinforce these small, randomly occurring clusters. Importantly, this global scenario is expected to require that the small synaptic clusters can control global synaptic plasticity by driving the output of the cell via the amplification of their postsynaptic effect by local dendritic nonlinearities.

In the present paper we develop a novel analysis method to estimate the effect of synaptic clustering on the post-synaptic response of a biophysical model neuron under *in vivo*-like conditions. Using our method we show that, when the connectivity is unstructured, small synaptic clusters do not influence the sVm response of CA1 and L2/3 pyramidal neuron models making the global model of cluster formation unfeasible. We further show that assuming uniform synaptic strength, 10-20 synapses per cluster are required to achieve reliable output tuning, but changing the strength or the number of inputs has a stronger effect on the neuronal output. Finally we show that the increase of the background activity during hippocampal sharp waves (SPW) paradoxically decreases the effect of synaptic clustering on the sVm.

## Results

### Clustering via global plasticity requires activation of dendritic nonlinearities

To examine the conditions for creating functional synaptic clusters via global mechanisms, we simulated structural plasticity in a simplified neuron model equipped with multiple nonlinear subunits, corresponding to idealised dendritic branches (Figure S1A-B). Importantly, in our simulations the structural synaptic plasticity was controlled by the output of the neuron. As expected, when subunit nonlinearities were weak, and thus synaptic clusters did not contribute to the variability of the neuronal output (Figure S1C-D), global plasticity did not lead to the formation of functional synaptic clusters (Figure S1F-G). However, when the effect of randomly occurring synaptic clusters was boosted by strong subunit nonlinearities (Figure S1B), and thus subunit nonlinearities contributed significantly to response variability (Figure S1E), the same global plasticity process led to the formation of functional synaptic clusters (Figure S1F-G).

These simulations demonstrated that functional clustering can be achieved via global plasticity mechanism when local nonlinearities contribute to the neuronal response variability. In the following section we investigate whether local dendritic nonlinearities in cortical pyramidal neurons are sufficiently strong to amplify the effect of small, randomly occurring input clusters and to control the neuronal responses under realistic input conditions.

### Randomly occuring synaptic clusters have small impact on neuronal responses

To estimate the impact of functional synaptic clustering on the somatic response under *in vivo*-like conditions we developed a novel analysis termed the *decomposition of response variance* (STAR Methods). In short, we simulated a biophysical model neuron whose integrative properties have been fitted to *in vitro* data, and stimulated it with input patterns matched to the input the neuron receives under *in vivo* conditions via excitatory and inhibitory synapses distributed throughout the entire dendritic tree. We recorded and analysed the biophysical model’s sVm response while manipulating the variability of the input and the fine-scale arrangement of the synapses.

We used a detailed model of a CA1 pyramidal cell (Jarsky et al. 2005; STAR Methods) that reproduced several somatic and dendritic properties of these cells measured under *in vitro* conditions (Figure S2), including the generation and propagation of Na^+^ action potentials at the soma and along the apical dendritic trunk (Jarsky et al., 2005); the generation of local Na^+^ spikes in thin dendritic branches (Losonczy & Magee, 2006); amplitude distribution of synaptic responses (Magee & Cook, 2000); nonlinear integration of inputs via NMDA receptors (NMDAR; Losonczy & Magee (2006)); the similar voltage threshold for Na^+^ and NMDA nonlinearities (Losonczy & Magee, 2006) and the major role of A-type K^+^ channels in limiting dendritic excitability (Hoffman et al., 1997; Losonczy & Magee, 2006).

After validating the biophysical model, we stimulated the neuron with synaptic input patterns characteristic for the hippocampal population activity under natural conditions (STAR Methods). Specifically, we simulated the activity of 2000 excitatory and 200 inhibitory presynaptic neurons during the movement of a mouse in a 2m long circular track (Figure 1A). The excitatory neurons exhibited a single idealised place field, were modulated by theta oscillation and showed phase precession (Figure S2H-I, Skaggs et al. 1996). As about 10% of hippocampal CA3 neurons show location selective activity in a given environment (Alme et al., 2014; Hainmueller & Bartos, 2018), the activity of the simulated cells faithfully represents the CA3 inputs received by the a postsynaptic CA1 neuron under *in vivo* conditions (STAR Methods). Inhibitory interneurons were modulated by theta oscillation (Csicsvári et al., 1999) but were spatially untuned (Dupret et al., 2013). Initially we chose unstructured connectivity between the inputs and the postsynaptic dendritic tree (*random synaptic arrangement*, Figure 1E). We studied the effects of small functional synaptic clusters (typically 2-4 synapses, depending on the definition of the synaptic cluster, Figure S3) occurring randomly under these conditions, on the sVm response of the neuron.

**Figure 1.**
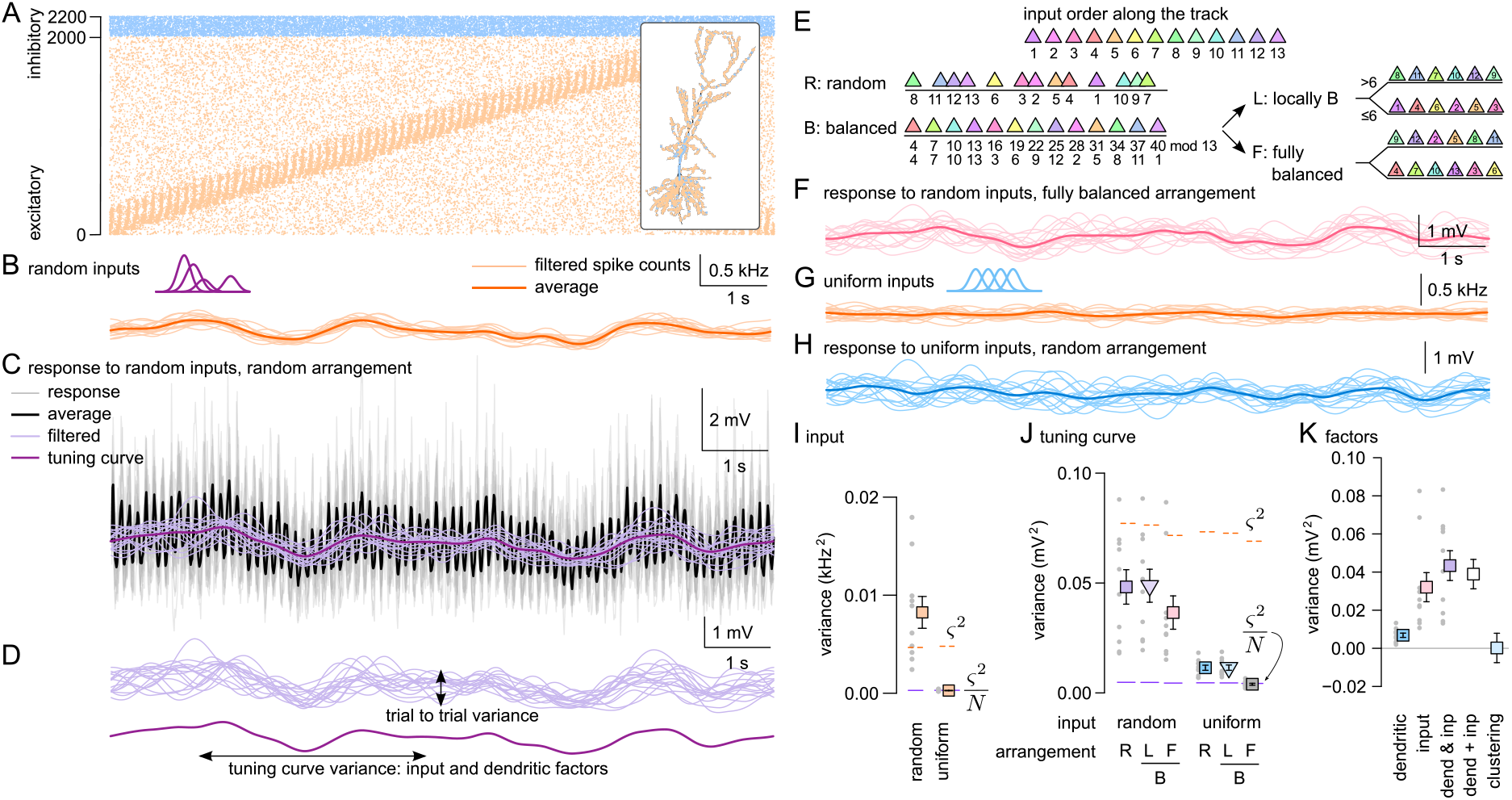
Random synaptic clusters have small impact on neuronal tuning. (A) Simulated activity of excitatory (orange) and inhibitory (blue) inputs in a single lap on the 2m long circular track. Excitatory inputs are ordered according to the location of their place field. The speed of the animal was constant 20 cm/s. Inset: Morphology of the modelled CA1 pyramidal neuron and the spatial distribution of the 2000 excitatory (orange) and 200 inhibitory (blue) synapses. (B) Filtered input spike counts in individual trials (light) and average (dark) in the random input condition (schematic in purple). (C) The sVm response of the postsynaptic CA1 pyramidal cell to inputs in 16 laps (gray) and the average postsynaptic response (black) in the random input condition with random synaptic arrangement. To remove theta modulation, we low-pass filtered the raw Vm traces (light purple). The average of the low-pass filtered traces yields the *tuning curve* (dark purple). (D) Illustration of the trial-to-trial variability, which is quantified by the average variance of the n=16 low-pass filtered responses (top) and the tuning curve variance which is the variance of the tuning curve along the track. (E) Schematic for the synaptic arrangements used in the paper. Top: ordering of 13 input place cells along the circular track. Color difference is proportional to place field distance. Left: random (R) and balanced (B) synaptic arrangement on a short segment of a single dendritic branch. Right: arranging inputs within each dendritic branch separately yields the locally balanced (L) arrangement that preserves large-scale inhomogeneities between branches (top, synapses 1-6 are on the lower branch) and fully balanced (F) arrangement that removes both small-scale clustering and large-scale inhomogeneities (bottom). (F) Filtered sVm response in individual trials (light) and average (dark) in the random input condition with fully balanced arrangement. (G) Filtered input spike counts in individual trials (light) and average (dark) in the uniform input condition (schematic in blue). (H) Filtered sVm response in individual trials (light) and average (dark) in the uniform input condition with random synaptic arrangement. (I) Variability of the inputs. The variance of the mean filtered spike count (shown in panel B and G) is higher than (same as) the variance expected from trial-to-trial variability in the random (uniform) input condition, respectively. Solid purple line segments show the variance expected based on trial-to-trial variability 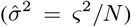 and dashed orange segments indicate trial-to-trial variability (*ς*^2^). (J) Tuning curve variance (as defined in panel F) in the random and uniform input conditions with random (R), locally balanced (F) and fully balanced (F) synaptic arrangements. (K) The contribution of the dendritic and input factors to the response variance. Grey dots in I-K show 10 independent simulations with different synaptic arrangements and inputs, symbols and error bars show mean and SEM.

We observed considerable variability of the sVm response both across trials and along the track (Figure 1C). To evaluate the contribution of synaptic clustering to the variability, we compared its effect to other components contributing to the total variance of the neuronal response. In particular, the sVm response variability can be attributed to three separate factors (Figure 1B-D): 1) trial-to-trial variability associated with stochastic biophysical processes (e.g., spiking and synaptic vesicle release) modeled as a Poisson process here (STAR Methods); 2) variability in the inputs active along the track, including variations in the precise number of presynaptic neurons with place fields at the current location, their maximal firing rates or their synaptic strengths; 3) dendritic factors, i.e., differential spatial distribution of inputs active along the track including small-scale functional clustering or large-scale spatial heterogeneities (e.g., proximal vs. distal dendritic location). Importantly, in our simulations we could separate the effect of the latter two factors by simple manipulations of the input conditions and the synaptic arrangement.

The trial-to-trial variability, *ς*^2^, was measured as the variance across *N* =16 trials with identical presynaptic firing rates but random presynaptic action potentials and synaptic release events (Figure 1C-D). Averaging the sVm across these trials provided the postsynaptic tuning curve along the track (Figure 1C-D). The trial-to-trial variability provided a natural lower bound on the expected variance of the tuning curve 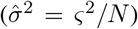. We first measured the contribution of the input and dendritic factors together on the variance of the tuning curve using random input conditions (Figure 1B, randomly varying the strength of the synapses and the location and amplitude of the presynaptic place fields within physiological range – STAR Methods), and random synaptic arrangement in 10 independent simulations. We found that the tuning curve varied considerably along the track (Figure 1C-D), and this tuning curve variability was consistently larger than the lower bound imposed by the trial-to-trial variability 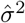 (Figure 1J).

Second, we eliminated the effect of dendritic factors on the response variability by rearranging synapses throughout the dendritic tree to minimize the correlation between the activity of nearby synapses (balanced synaptic arrangement, Figure 1E, Figure S4 and STAR Methods). In this balanced arrangement the functional synaptic clusters are equally absent for all differently tuned presynaptic ensembles, while dendritic processing in general is maintained. In the first step we rearranged synapses only within individual dendritic branches, removing local synaptic clusters but otherwise keeping large-scale, global biases intact (locally balanced synaptic arrangement, L). We found that this manipulation did not change the response variance compared to the random synaptic arrangement (P=0.99; Kolmogorov-Smirnov test, Figure 1J), indicating that small functional clusters occurring randomly had no effect on the neuronal tuning. Next, we fully rearranged the synapses removing both local clusters and large-scale inhomogeneities (*fully balanced* synaptic arrangement). This manipulation slightly decreased the tuning curve variance, but the effect was not significant because of the large variability across simulations (Figure 1F, J, random vs. fully balanced arrangement, P=0.45, KS-test).

Third, to isolate and directly measure the effect of dendritic factors, we eliminated the impact of input factors by setting the strength of the synapses identical and setting the total input rate to constant, but keeping the inputs otherwise unchanged. To achieve this we fixed the shape of the place fields and organized them to uniformly tile the environment (*‘uniform’ input condition*, Figure 1G). This way, the variability of the input was minimised (Figure 1I) and thus all remaining variability in the tuning curve was attributed to dendritic factors (Figure S3). Using random synaptic arrangement, we found that after eliminating the input factors the variability of the tuning curve decreased substantially but the remaining variability was still larger than 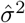 (Figure 1H, J, uniform vs. random input condition, P=0.0002, KS-test). We also observed large reduction in the variability between the 10 simulated neurons indicating that the majority of the cell-to-cell variability was caused by the random selection of inputs. Rearranging synapses within individual dendritic branches did not have an effect on the tuning curve variability (Figure 1J, uniform input, random vs. locally balanced synaptic arrangement, P=0.79, KS-test). Finally, when uniform input condition was combined with fully balanced synaptic arrangement, the tuning curve variance became practically identical to the variance expected from the trial-to-trial variability (Figure 1J, uniform input, balanced arrangement vs. 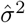, P=0.052, KS-test), confirming that our manipulations successfully eliminated both input and dendritic factors. We obtained similar result with L2/3 pyramidal neurons using inputs matched to population activity recorded in the visual cortex *in vivo* (Figure S5A-E).

From these results we conclude that with unstructured connectivity, input factors provide a substantially stronger contribution to the sVm response of CA1 and L2/3 pyramidal cells than dendritic factors. Moreover, we found that among dendritic factors, large-scale biases in the synaptic input tuning have a measurable contribution to the response variability, but small functional synaptic clusters do not influence neuronal responses (Figure 1K). As the output of the neuron is controlled by these other factors, back-propagating action potentials are not correlated with the activity of the small synaptic clusters and thus, synaptic clusters can not be reliably reinforced via global, Hebbian plasticity (Figure S1). Our data therefore supports the need of local synaptic plasticity mechanisms with low activation threshold for the formation of synaptic clusters (Weber et al., 2016).

Next we introduce structured synaptic connectivity (synaptic clusters) and study how they influence the tuning curve of the neuron.

### Larger synaptic clusters can lead to clustering-based tuning

To systematically study the effect of synaptic clustering we chose a subset of excitatory synapses that were coactive in the middle of the track (Figure 2A) and organised them into clusters of increasing size (Figure 2B; STAR Methods). Specifically, we increased the cluster size from 1 synapse per cluster (no clustering) to 60 synapses per cluster (maximal cluster size) in 7 discrete steps, and measured the sVm response of the neuron with different levels of clustering to identical presynaptic inputs (Figure 2C). While we did not observe a detectable effect of clusters consisting of 2-5 synapses, when cluster size reached that of 10 synapses, a depolarization ramp emerged in the tuning curve at the location where the clustered synapses were active (Figure 2D) increasing the variance of the tuning curve (Figure 2E) and the response integral (Figure 2F). Note that the total synaptic input the neuron receives does not change along the track (‘uniform’ input condition) and the ramp is the somatic signature of nonlinear dendritic processing of clustered inputs: the tuning is entirely due to input clustering. Including dendritic spines in the simulations did not drastically change the minimal cluster size required to achieve reliable tuning (Figure 2E-F). Further increasing the cluster size to 20 synapses per cluster increased the magnitude of the depolarizing ramp, but it also saturated the response (Figure 2E-F).

**Figure 2.**
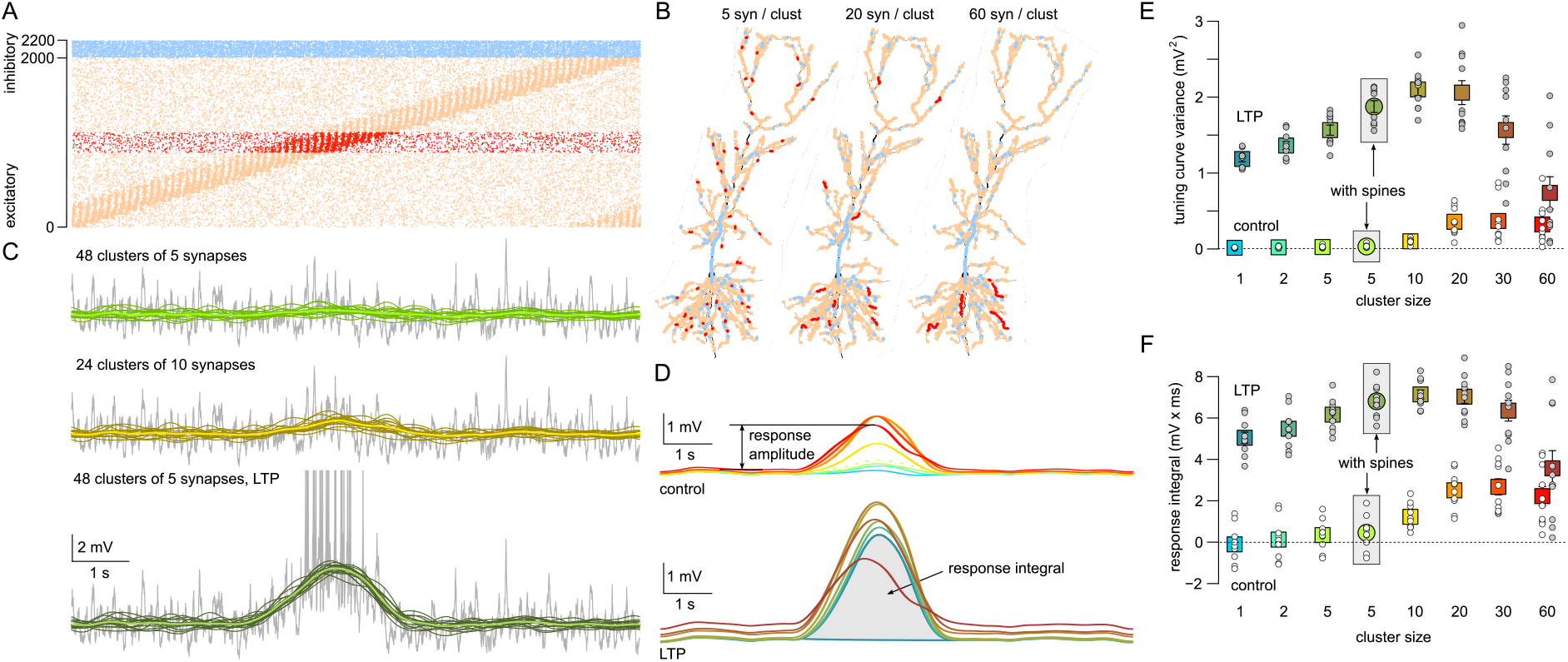
Larger synaptic clusters can lead to clustering based tuning. (A) Excitatory inputs coactive in the middle of the track (red) were chosen for clustering. (B) Branches longer than 60 *μ*m were selected for arranging 240 synapses (red) into 48 (left), 12 (middle) and 4 (right) functional synaptic clusters. Background excitatory (orange) and inhibitory (blue) synapses are also shown. (C) Example sVm response of the CA1 neuron (grey), filtered responses for 16 trials (green and olive) and tuning curve (light green and yellow) with 5 (top) or 10 synapses per cluster (middle) or with 5 synapses per cluster combined with LTP of the clustered synapses. Spikes were removed from the responses to obtain the filtered responses and the tuning curve. (D) Average subthreshold response of the CA1 postsynaptic cell shows increased depolarization ramp amplitude with clustering (top). Potentiation of the clustered synapses further increases the ramp amplitude (bottom). Color code is the same as in panels E and F. (E) Tuning curve variance as a function of synaptic clustering for control (bright colors) and LTP (dark colors) in 10 independent simulations (circles) and their means (color boxes) and SEM (error bars). Data from simulations including dendritic spines and 5 cluster per branches is highlighted by grey boxes. Note that the placements of the red synapses is biased in towards long terminal branches even in the absence of clustering. (F) Tuning curve ramp integral as a function of synaptic clustering for control (bright colors) and LTP (dark colors) in 10 independent simulations (circles) and their means (color boxes) and SEM (error bars).

As synaptic factors play an important role in establishing the tuning of hippocampal place cells (Bittner et al., 2017), we compared the effect of synaptic clustering on the tuning curve with the possible effects of synaptic plasticity. As a simple model of synaptic changes during LTP, we doubled the conductance of the clustered synapses and recorded the tuning curve while changing the level of clustering. We found that the effect of synaptic plasticity was larger than the effect achieved by clustering in general (Figure 2C-F). Similar to the control case, maximal depolarization of the tuning curve was achieved at intermediate cluster sizes (10-20 synapse per cluster, Figure 2D) but further increasing the level of clustering severely saturated the membrane potential of dendritic branches receiving clustered inputs, and decreased the average magnitude of NMDAR current per synapse (Figure 3D). We obtained similar results in the L2/3 neuron: clusters of 10-20 synapses lead to maximal responses and larger clusters yielded saturation (Figure S5F-J). Furthermore, we found in the CA1 neuron model that dendritic spines increased the influence of small synaptic clusters when the clustered synapses were potentiated, but had little effect on the responses without potentiation (Figure 2E-F).

**Figure 3.**
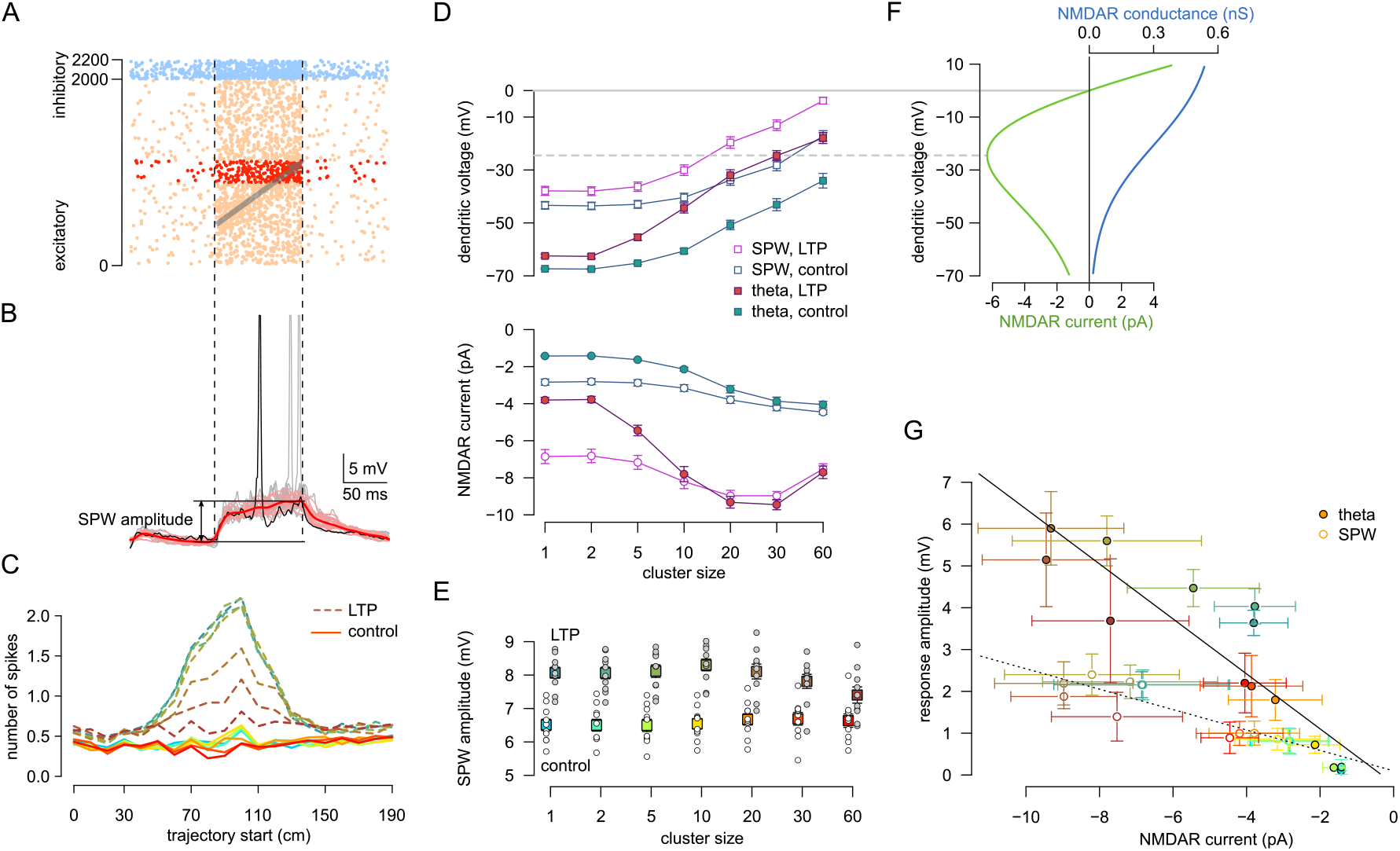
Synaptic clustering have reduced impact during elevated states. (A) Simulated inhibitory (blue) and excitatory (orange and red) inputs during an example SPW (vertical dashed lines). Grey arrow indicates the replayed trajectory. Red dots show activity of the synapses arranged into functional clusters (see Figure 2B). Note that the high background firing also includes the activity of input neurons not having a place field on this particular track. (B) Postsynaptic membrane potential response to SPW inputs. Black trace: response to the inputs in panel A; grey: 15 repetitions with the same presynaptic firing rate but random inputs; pink: subthreshold, filtered response; red: average postsynaptic response, which was used to measure the amplitude of the depolarisation relative to the pre-SPW period. (C) Average number of spikes during individual SPW events as a function of the start location of the trajectory for control (light colors) and LTP (dashed lines). Color code: same as in panel E. The largest firing rate is when the trajectory starts at 100 cm, which is analysed further in panels D-G in the case of passive neuron. (D) Local dendritic voltage at the synaptic clusters (top) and average NMDAR current of clustered synapses (bottom) during the peak activity of the clustered synapses in SPW events (open symbols) and during theta state (filled symbols). Error bars indicate standard error of the mean across 40 dendritic branches (top) or 40 synapses (bottom). (E) Mean somatic depolarisation amplitude relative to the pre-SPW baseline (B) as a function of cluster size for control (bright colors) and LTP (dark colors, trajectory start =100 cm). Circles show 10 independent simulations with different cluster arrangements and inputs; symbols and error bars show mean and SEM. (F) NMDAR conductance (blue) and current (green) as a function of voltage. The peak NMDAR current is at ~-25 mV (dashed grey line). (G) Somatic response amplitude as a function of the average NMDAR current during theta (filled circles) and SPWs (open circles). Error bars show standard deviation across 10 simulations (vertical) or 40 synapses (horizontal). The high variability is explained by the large diversity between dendritic branches receiving clustered inputs. Here the response amplitude during SPWs is measured relative to the depolarisation during the replay of trajectories that do not involve clustered synapses. The response amplitude during theta is the maximum of the average subthreshold response minus the baseline (Figure 2E).

Our data shows that during place cell activity and theta oscillation, clustering-based tuning emerges in CA1 pyramidal cells only if the clusters are relatively large. However, neuronal signal integration is highly dependent on the state of the network (Destexhe et al., 2003). Therefore in the next section we examined the effect of synaptic clustering on the postsynaptic response during hippocampal sharp waves.

### Synaptic clustering has small impact during hippocampal sharp waves

Hippocampal sharp waves (SPW) are transient, highly synchronous network states where a large fraction of the local network becomes active (Buzsáki, 2015). During SPWs, the population activity internally recreates sequential activity patterns experienced earlier during exploratory behavior and theta activity (Foster, 2017). We expected the critical cluster size to be smaller during SPWs than during theta activity, as elevated excitatory activity can facilitate the generation of dendritic spikes (Farinella et al., 2014).

We simulated replay events embedded in an elevated background activity, mimicking hippocampal population activity during SPW-ripples (Figure 3A). The statistics of the excitatory and inhibitory inputs, including the population firing rates, the ripple modulation of the cells and the statistics of the replay events were matched to *in vivo* data (Mizuseki & Buzsáki 2013; Pfeiffer & Foster 2013; English et al. 2014, STAR Methods). We systematically varied the overlap between the replayed trajectory with the place fields of the neurons giving postsynaptically clustered inputs (Figure 3A-B). Contrary to our expectations, we found that the number of spikes during individual sharp wave events (Figure 3C) and the amplitude of the somatic depolarisation (Figure 3E) were insensitive of the arrangement of the synapses into functional clusters. Increasing the conductance of the selected excitatory synapses by a factor of 2 (mimicking LTP) resulted in a substantially larger output spike count and somatic depolarisation amplitude when clusters were small (Figure 3C, E). However, when LTP was combined with large functional synaptic clusters the advantage of the LTP began to disappear and the output spike counts and somatic depolarisation amplitudes were reduced (Figure 3C, E).

To understand the biophysical mechanisms underlying the paradoxical effects of functional clustering during SPWs, we analysed how the local dendritic depolarization and the average NMDAR current of a single synapse within a functional cluster changes with the size of the cluster (Figure 3D). We could identify two different mechanisms that contributed to the reduced impact of clustering during sharp waves. First, local dendritic voltage was substantially more depolarised during SPWs than during theta (Figure 3D, top). Although clustering further increased the local depolarisation during SPWs, it had only a minor impact on the average NMDAR current per synapse (Figure 3D, bottom) as that was already near-maximal even in the absence of clustering (cluster size = 1; Figure 3F). Second, the elevated synaptic conductance load during SPWs reduced the gain of the neuronal responses such that similar changes in the input currents had a markedly smaller effect on the postsynaptic responses (Figure 3G, Mel 1993).

These results demonstrate that the impact of synaptic clustering on the neural response is small during SPWs and can even be negative when large clusters are combined with LTP and synaptic saturation becomes the dominant effect.

## Discussion

### Contribution of Dendritic and Synaptic factors to Neuronal Tuning

Identifying the external sensory and internal biophysical correlates of neuronal tuning has been a longstanding goal in systems neuroscience (Rieke et al., 1996; Wilson et al., 2018). Previous approaches to estimates of the contribution of dendritic nonlinearities to stimulus selectivity relied on pharmacological manipulations which also influenced synaptic factors (Smith et al., 2013; Palmer et al., 2014; Lavzin et al., 2012). Our computational approach offers a novel way to isolate these effects and measure the contribution of dendritic and synaptic factors on the neuronal tuning and compare it to the trial-to-trial variability.

Our conclusions assume that we could accurately estimate the magnitude of these factors. In particular, severe underestimation of dendritic and overestimation of synaptic factors could have a major impact on the conclusions. To ensure that these factors are correctly estimated, we matched the synaptic conductances (Nicholson et al., 2006), the EPSP amplitudes (Magee & Cook, 2000) and the dendritic active conductances to experimental data (Jarsky et al., 2005; Losonczy & Magee, 2006). Stronger dendritic nonlinearities, such as fully propagating dendritic Na+ spikes in hippocampal interneurons (Martina et al., 2000) or bistable NMDAR nonlinearities (Major et al., 2013; Schiller et al., 2000) in L5 neurons could increase the contribution of dendritic factors. Conversely, other factors, such as larger differences in the place field properties, greater variability in the synaptic receptor numbers (Nusser et al., 1998) or correlations between the presynaptic firing rates and the synaptic strength (Koulakov et al., 2009) are expected to increase the influence of the synaptic factors. In addition, state-dependent changes in neuromodulation may dynamically regulate the relative contribution of synaptic inputs and dendritic processing *in vivo* (Williams & Fletcher, 2019).

Local dendritic nonlinearities, in principle, can fundamentally change the integrative properties of neurons and can also have a profound effect on the computational capabilities of the circuit. However, such a large computational effect requires very strong local, branch-specific nonlinearities (e.g., Poirazi & Mel 2001 used *g*(*x*) = *x*^10^ as a subunit nonlinearity) and a large influence on the global output. This is in contrast with our findings that the effect of local dendritic nonlinearities on the somatic response of the neuron under *in vivo*-like conditions is relatively small compared to synaptic factors or global, neuron-wide nonlinearities at least in CA1 and L2/3 pyramidal cells (Ujfalussy et al., 2018).

### Local vs. global plasticity

Using a simple model we demonstrated that functional synaptic clusters can be formed and stabilized via global synaptic plasticity rules only in the presence of strong local dendritic nonlinearities (Figure S1). This mechanism did not generate clusters composed of functionally homogenous inputs, as global plasticity stabilises any input irrespective of its location within the dendritic tree that show correlated activity with a functional clusters. The interspersion of inputs with a wide range of functional properties on individual branches is an important characteristic of experimental data (Jia et al., 2010; Scholl et al., 2017; Wilson et al., 2018) and may thus be a signature of the regulation of structural synaptic plasticity by global factors. Although the functional clustering observed in the model was statistically significant, the magnitude of the effect (difference between the observed and the random) was not large. This is, again, reminiscent of the experimental data, where the difference between the data and the appropriately shuffled control distribution is often relatively small (Takahashi et al., 2012; Iacaruso et al., 2017; Scholl et al., 2017; Kerlin et al., 2018). Importantly, to generate functional clusters by global plasticity alone, dendritic nonlinearities have to be sufficiently strong (Figure S1) and should have a low threshold to amplify the effect of small, randomly occurring clusters and to control the plasticity process, whereas clustering via local plasticity mechanisms does not necessarily require dendritic nonlinearities (Weber et al., 2016). We found that synaptic clusters randomly occurring within dendritic branches did not contribute to the tuning curve variability and thus global plasticity per se is unlikely to account for the reinforcement of small functional synaptic clusters. However, global plasticity can reinforce large scale biases in synaptic tuning properties (Iacaruso et al., 2017) as these factors did have a measurable contribution to the neuronal tuning curve. The interactions between local and global factors in controlling synaptic plasticity and the generation of functional clusters require further investigations.

### Cluster size

Under *in vitro* conditions the induction of dendritic spikes requires the near-synchron activation of a sufficiently large number of inputs (Schiller et al., 2000; Losonczy & Magee, 2006; Branco & Häusser, 2011; Makara & Magee, 2013), but whether inputs occurring *in vivo* have the sufficient synchrony to trigger them is debated (London et al., 2010; Yuste, 2011). Here we simulated *in* vivo-like activity of a full presynaptic population carefully matched to experimental data to ensure that we capture the correlations present in the inputs to pyramidal neurons. We found that dendritic nonlinearities could be activated when the size of the functional clusters was comparable to the number of stimuli eliciting dendritic spikes *in vitro*. Although the spatial scale of the functional clusters found in cortical neurons has been restricted to 5-10 *μ*m and 2-5 dendritic spines (Takahashi et al., 2012; Iacaruso et al., 2017; Scholl et al., 2017), current experimental techniques do not allow reliable monitoring the activity of all synaptic inputs in a given dendritic branch and thus, these studies could easily underestimate the real number of synapses involved in a given synaptic cluster.

The current flowing through an NMDA receptor depends on the local voltage, which is the function of the activity of the neighbouring synapses. The optimal level of synaptic clustering for a given input statistics to effect the somatic response can be defined as the arrangement of the inputs that maximises the NMDAR current per synapse (Mel, 1993). There are several important points to note in this definition. First, the optimal cluster size to influence somatic tuning depends on the input: for larger input firing rates smaller clusters will be optimal - as we observed in the case of sharp waves and theta. Second, optimal cluster size is different for each dendritic branch, as smaller clusters will achieve larger depolarisation in branches with higher input resistance. Finally, the optimality of the cluster size in a given branch can be tested experimentally by recording the membrane potential distribution of dendritic branches *in vivo* (Smith et al., 2013) and calculating the mean NMDAR current.

### Computational role of synaptic clustering

The prevailing computational theory of local dendritic nonlinearities states that they increase the representational capacity of single neurons by allowing them to correctly classify inputs that are linearly not separable (Poirazi & Mel, 2001; Schiess et al., 2016). Because in this computation the mapping between inputs and dendritic branches is determined by the target classification rule and not by the correlations present in the inputs, this computation inherently predicts that the formation of synaptic clusters targeting dendritic subunits is regulated globally. On the other hand, functional clustering based on input correlations has been predicted by other theoretical considerations, such as robustness to noise (Cazé et al., 2017) or efficient spike-based computations (Ujfalussy et al., 2015). In particular, this latter theory also predicts a match between the type and magnitude of input correlations and the form of dendritic integration which is independent of the global neuronal responses and thus has to be regulated by local dendritic plasticity rules.

The temporal window for interaction between inputs targeting neighbouring dendritic spines for synaptic plasticity was often found to be orders of magnitude larger (≈ 10 min, Harvey & Svoboda 2007; Govindarajan et al. 2011) than the synchrony required for nonlinear dendritic integration (< 10 ms, Losonczy & Magee 2006). Even if the dendritic nonlinearities are too weak to substantially amplify the neuronal responses to clustered inputs or inputs are too asynchronous to trigger dendritic spikes, synaptic clustering could have an important role in reducing the interference between memories and to promote selective generalisation via spatially restricting the effect of plasticity-related molecules (Harvey & Svoboda, 2007; Govindarajan et al., 2011; O’Donnell & Sejnowski, 2014).

In conclusion, our findings indicate that 1) the selective responses of cortical neurons are primarily the consequence of the tuning of their synaptic inputs, 2) functional synaptic clustering matched to local dendritic properties can have additional role in refining those responses, 3) plasticity of functional synapse clusters such as those observed *in vivo* requires local rather than global mechanisms, and 4) in turn, local plasticity by small synaptic clusters may lead to powerful tuning of somatic responses.

## Acknowledgements

This work was supported by an MTA and an NKFIH fellowship (PD-020/2015 and PD-125386, FK-125324; B.B.U.), the Howard Hughes Medical Institute (55008740; J.K.M.), and the ERC (CoG 771849; J.K.M.). We thank Lajos Vágó for the idea of co-prime ordering and Zoltán Nusser for discussions and for his comments on an earlier version of the manuscript.

## Author Contributions

B.B.U., J.K.M. designed the study. B.B.U. performed the simulations and analyzed the data. J.K.M. and B.B.U. interpreted the results and wrote the manuscript.

## STAR Methods

### KEY RESOURCES TABLE CONTACT FOR REAGENT AND RESOURCE SHARING

As Lead Contact, Balázs B Ujfalussy is responsible for all reagent and resource requests. Please contact Balázs B Ujfalussy at with requests and inquiries.

### METHOD DETAILS

#### Biophysical models

Simulations were performed with the NEURON simulation environment (Hines & Carnevale 1997 version 7.4) embedded in Python 2.7 (Rossum, 1995).

##### CA1 neuron model

We used a modified version of the CA1 pyramidal cell model of Jarsky et al. (2005) to better account for the dendritic processing of synaptic inputs in CA1 pyramidal neurons (Losonczy & Magee, 2006). The passive parameters of the model were slightly adjusted to capture the dendro-somatic attenuation of synaptic responses: *C*_m_ = 1 *μ*F/cm^2^, *R*_i_ = 100 Ωcm and *R*_m_ = 20kΩcm^2^ in the dendrites, *R*_m_ = 40kΩcm^2^ in the soma and in the axon and *R*_m_ = 50 Ωcm^2^ in the axonal nodes. To correct for the presence of spines, *C*_m_ was increased and *R*_m_ was decreased by a factor of 2 in dendritic compartments beyond 100 *μ*m from the soma. In some simulations the excitatory synapses were placed on dendritic spines consisting of a spine neck (length: 1.58 *μ*m, diameter: 0.077 *μ*m) and spine head (length: 0.5 *μ*m, diameter: 0.5 *μ*m) with total neck resistance ~500 MΩ (Harnett et al., 2012). Since even in these simulations we only included about 10% of the spines present in CA1 pyramidal cells explicitly, we did not change the compensation factor.

To replicate important features of dendritic integration of excitatory synaptic inputs in the model we had to slightly modify the original ion channel parameters. We focused on local nonlinearities (i.e. Na^+^ and NMDA spikes) as we assumed that such spikes are more likely to be engaged by small synaptic clusters than global regenerative demdritic spikes such as plateau potentials (Larkum et al., 2009). Specifically, in order to increase the threshold for dendritic Na^+^ spike initiation to a similar value as the threshold for local NMDA spikes (Losonczy & Magee, 2006), the activation curve of the Na^+^ channels was shifted by 20mV in the basal, oblique and tuft branches (but not in the apical trunk). Furthermore, to prevent the attenuation of NMDAR currents by the activation of K^+^channels after local Na^+^ spikes, we decreased the density of the K^+^ channels in the dendritic branches (Table 1). Finally, in the original model (Jarsky et al., 2005) action potentials were initialised in the axonal nodes and propagated actively to the soma. Under *in vivo*-like synaptic inputs, the high conductance load at the soma often prevented the generation of full action potentials in the original model and in these cases axonal spikes appeared as spikelets (Michalikova et al., 2017). To eliminate spikelets (Hulse et al., 2016) we increased the somatic and decreased the axonal Na+channel conductance (Table 1).

**Table 1.** Ion channel densities in the model. The distance along the trunk, *d*, is measured in *μ*m. As in Jarsky et al. (2005), the dendritic segments closer than 100 *μ*m to the soma contained 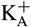 channels with lower half-inactivation voltage (Hoffman et al., 1997).

| | $\text{Na}^+ (\text{S}/\text{cm}^2)$ | $\text{K}_{\text{DR}}^+ (\text{S}/\text{cm}^2)$ | $\text{K}_A^+ (\text{S}/\text{cm}^2)$ |
| --- | --- | --- | --- |
| apical trunk ($< 100 \mu\text{m}$) | 0.04 | 0.04 | $\min\{0.048 (1 + d/100), 0.288\}$ |
| apical trunk ($> 100 \mu\text{m}$) | $\min\{0.04 (1 + d/1000), 0.06\}$ | 0.04 | $\min\{0.048 (1 + d/100), 0.288\}$ |
| other dendrites | 0.03 | 0.02 | 0.02 |
| soma | 0.2 | 0.04 | 0.02 |
| axon hillock | 0.04 | 0.04 | 0 |
| axon initial segment | 0.04 | 0.04 | 0.02 |
| axonal inter-nodal segment | 0.04 | 0.04 | 0.004 |
| axonal nodes | 15 | 0.04 | 0.004 |

When we measured the dendritic voltage and NMDAR currents in Figure 3D-G, we used a passive version of the model with all active conductances set to zero except the conductance of NMDA receptors.

##### L2/3 neuron model

For the L2/3 pyramidal neuron model shown in Figure S5 we used a detailed reconstruction of a biocytin-filled layer 2/3 pyramidal neuron (NeuroMorpho.org ID Martin, NMO-00904) as described previously (Smith et al., 2013; Ujfalussy et al., 2018). The passive parameters were *C*_m_ = 1 *μ*F/cm^2^, *R*_m_ = 7000 Ωcm^2^, *R*_i_ = 100Ωcm, yielding a somatic input resistance of 70MΩ.

Active conductances were added to all dendritic compartments and to the soma and included the following: voltage-activated Na^+^ channels (soma 100 mS/cm^2^, dendrite 8 mS/cm^2^ and hot spots 60 mS/cm^2^, Nevian et al. 2007); voltage-activated K^+^ channels (10 mS/cm^2^ soma and 0.3 mS/cm^2^ dendrite); M-type K^+^ channels (soma 0.22 mS/cm^2^ and dendrite 0.1 mS/cm^2^); Ca^2+^-activated K^+^ channels (soma 0.3 mS/cm^2^ and dendrite 0.3 mS/cm^2^); high-voltage activated Ca^2+^ channels (soma 0.05 mS/cm^2^ and dendrite 0.05 mS/cm^2^) and low-voltage activated Ca^2+^ channels (soma 0.3 mS/cm^2^ and dendrite 0.15mS/cm^2^). Calcium handling was modeled by a first-order system representing Ca^2+^ pumps and buffers with a time constant of decay of Ca^2+^ of 28.6 ms and the equilibrium free intracellular Ca^2+^ concentration of 100 nM.

##### Synapses

The model included AMPA and NMDA excitation and slow and fast GABAergic inhibition with synaptic conductances modeled as double-exponential functions. Each excitatory synapse included an AMPA and a NMDA component which were thus colocalized and always coactivated. Similarly, inhibitory synapses were composed of a mixture of GABA-A and GABA-B receptors. The parameters of the synaptic conductances are shown in Table 2 for both the L2/3 and the CA1 neuron model. The voltage dependence of the NMDAR conductance was captured by the standard sigmoidal activation curve:

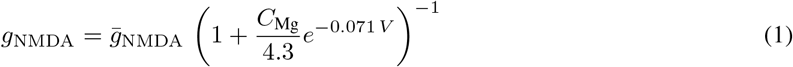

with the Mg^2+^ concentration beeing *C*_Mg_ = 1 mM and with a slightly steeper activation than in the original model Jahr & Stevens (1993). The maximal conductance of both the NMDA and AMPA synapses was doubled for the clustered synapses when we modeled LTP.

**Table 2.** Synaptic parameters used in the models. The maximal conductance of both the AMPA (NMDA) synapses was chosen randomly from a uniform distribution between 0.15 (0.2) and 1.05 (1.4) nS for the CA1 model during random input conditions, respectively.

|  |  | CA1 model | L2/3 model |
| --- | --- | --- | --- |
| AMPA | $\tau_1$ | $0.1 \text{ ms}$ | $0.1 \text{ ms}$ |
| | $\tau_2$ | $1 \text{ ms}$ | $2 \text{ ms}$ |
| | $g_{\text{max}}$ | $0.6 \text{ nS}$ | $0.5 \text{ nS}$ |
| | $E_{\text{rev}}$ | $0 \text{ mV}$ | $0 \text{ mV}$ |
| NMDA | $\tau_1$ | $2 \text{ ms}$ | $2 \text{ ms}$ |
| | $\tau_2$ | $50 \text{ ms}$ | $40 \text{ ms}$ |
| | $g_{\text{max}}$ | $0.8 \text{ nS}$ | $0.8 \text{ nS}$ |
| | $E_{\text{rev}}$ | $0 \text{ mV}$ | $0 \text{ mV}$ |
| $\text{GABA}_{\text{fast}}$ | $\tau_1$ | $0.1 \text{ ms}$ | $0.1 \text{ ms}$ |
| | $\tau_2$ | $4 \text{ ms}$ | $4 \text{ ms}$ |
| | $g_{\text{max}}$ | $0.7 \text{ nS}$ | $0.7 \text{ nS}$ |
| | $E_{\text{rev}}$ | $-65 \text{ mV}$ | $-70 \text{ mV}$ |
| $\text{GABA}_{\text{slow}}$ | $\tau_1$ | $1 \text{ ms}$ | $1 \text{ ms}$ |
| | $\tau_2$ | $40 \text{ ms}$ | $40 \text{ ms}$ |
| | $g_{\text{max}}$ | $1.2 \text{ nS}$ | $0.33 \text{ nS}$ |
| | $E_{\text{rev}}$ | $-80 \text{ mV}$ | $-85 \text{ mV}$ |

We simulated *N*_E_ = 2000 excitatory and *N*_I_ = 200 inhibitory synapses in the CA1 cell and *N*_E_ = 1920 excitatory and *N*_I_ = 192 inhibitory synapses in the L2/3 cell. In the random synaptic arrangement excitatory synapses were placed randomly with a uniform distribution on the entire dendritic tree of the postsynaptic neuron (Figure S4A).

When we systematically varied the size of the clusters, we selected 240 presynaptic inputs based on the location of their place field (or orientation tuning in L2/3 cell) and varied the arrangement of the corresponding synapses on the postsynaptic dendritic tree. Specifically, the inputs were ordered by the location of their place fields (orientation preferences) and were divided into *N*_clust_ ∈ {240, 120, 48, 24, 12, 8, 4} contiguous and disjoint sets of clusters, each containing *M*_clust_ ∈ {1, 2, 5, 10, 20, 30, 60} inputs. The remaining 1760 background inputs, having tuning curves negatively correlated with the tuning of the clusters, were placed randomly with uniform density. Clusters were placed on dendritic branches longer than *L* = 60 *μ*m by first randomly selecting a branch (with probability proportional to its length) and then a cluster starting point between the proximal end of the branch and *L* – *d M*_clust_ *μ*m distance from it. Synapses were added distally from the cluster starting point with inter-synapse distance of *d* = 1 *μ*m. This procedure guaranteed that synapses within one cluster target the same dendritic branch, show maximally correlated activity and that the expected location of clustered synapses is independent of the cluster size. Note, that selection of long branches, typically thin terminal dendrites, for the location of clustered synapses introduces a slight bias for the postsynaptic processing of clustered inputs even when *M*_clust_ = 1. This bias is responsible for the difference between Figure 1J and Figure 2E, *N*_clust_ = 1.

Also note, that branches that received clustered inputs had higher synapse density due to the presence of background inputs. This was not a problem during theta as the background activity is relatively low and asynchron in that case, but could be a substantial concern during sharp waves. To exclude the possibility that synaptic saturation during SPWs is caused by the increased inputs at clustered dendrites, we generated 2000 background input locations (instead of 1760) distributed uniformly and selected the *N*_syn_ = 240/*N*_clust_ background inputs closest to the location of the 240 clustered synapses. These background inputs did not participate in the replay of any particular trajectory (see below) but only fired at elevated baseline rate during SPWs. Conversely, the clustered synapses did not show an elevated baseline activity during SPWs but only participated in the trajectory replay. Using this procedure we achieved a similar average input level for branches receiving clustered versus non-clustered inputs. The data presented in Figure 3 used this more uniform input distribution, but similar results were achieved when clusters were simply added to the background (data not shown).

Inhibitory inputs were divided into two groups with 80 synapses targeting the soma (and the apical trunk in the case of the CA1 neuron) and the remaining synapses were distributed randomly along the entire dendritic tree.

##### Balanced synaptic arrangement

To achieve a balanced synaptic configuration (B in Figure 1I) in the case of the hippocampal pyramidal cell, we first applied *co-prime reordering* procedure on the *N* = 40 presynaptic ensembles (see below) that minimized correlation between the neighbouring ensembles. To achieve this, we selected base numbers *α* relative prime to *N* and generated an arithmetic progression with difference *α*, dividing the elements of the sequence by N and taking the remainder (Figure 1E). In the case of *α* = 3 and *N* = 10 a potential sequence is:

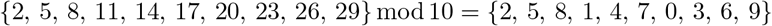

In this sequence the distance between neighbouring elements is constant, so input correlations are identical within each subset of the sequence. Also note that the sequence is not repeating until *N* elements. We chose *α* = 9 that minimized correlation between inputs within ~40 *μ*m (9 synapses).

Next, we arranged the 2000 inputs into a single sequence where cells from the different ensembles were selected according to the ordering defined above. Finally, we mapped this sequence to the dendritic tree of the neuron starting from the soma and following each subtree towards the distal end sequentially. This defined the fully balanced (F) arrangement (Figure S4D).

In the locally balanced, globally random arrangement (L, Figure 1E) we started with a random configuration and applied the co-prime reordering procedure independently to synapses targeting the same dendritic branch. Specifically, for *N_i_* synapses targeting branch *i* we selected *α_i_* ≈ *N_i_*/4 and rearranged the inputs within branch *i* while also equalizing the distance between neighbouring synapses (Figure S4C).

In addition, we also tested a globally balanced, locally random arrangement, where we started from the fully balanced configuration, and randomized synapse location and the ordering of the synapses within each branch separately (Figure S4B). The results achieved with this configuration were similar to that obtained with the fully balanced configuration (data not shown).

Visual cortical inputs targeting the L2/3 neuron model had 4 different features: orientation preference, phase preference, orientation selectivity and response linearity (Niell & Stryker, 2008). We co-sorted inputs to achieve an input distribution that is approximately balanced with respect to all 4 features at the same time (for details, see legend of Figure S4E-H).

#### Inputs

To study the impact of synaptic clustering under *in vivo*-like conditions in the biophysical models we carefully matched the inputs of the models to the synaptic inputs experienced by these neurons *in vivo*. Inputs, corresponding to synaptic release events, were generated by a Generalised Linear Model (GLM) tuned to a number of different external and internal features (Pillow, 2007; Chadwick et al., 2015). Specifically, the presynaptic input count at time *t* for synapse *i* were generated from a Poisson process with rate 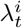:

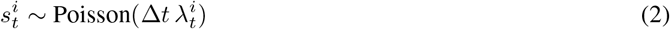

where the rate was defined as

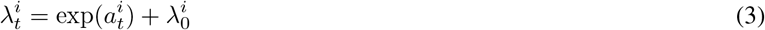

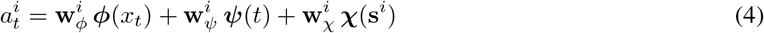

where 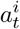 is called the activation and *ϕ*(*χ_t_*), *ψ*(*t*) and *χ*(s^*i*^) denote the activation of basis functions tuned to external inputs (location or stimulus orientation and phase), internal processes (phase of theta or gamma oscillation) or the output of the particular presynaptic neuron, respectively. The tuning properties of a particular input is set by the nature of these basis functions, varied among different conditions (theta or SPW for CA1 and gratings for L2/3) and parameter vector w^*i*^ specific for each cell *i*. The term 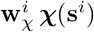 captures the effect of past events on the input rate (Pillow, 2007) and was included to model a short (5 ms) refractory period in the inputs.

Although the rate of each input was a deterministic function of time, the synaptic events were generated by a stochastic process accounting for variability in spike timing and stochastic vesicle release. This allowed us to generate multiple trials with identical input rates to estimate the trial-to-trial variability. Although the framework is identical for the three different conditions, we describe our choice for the basis functions and the parameters separately in the following sections. Inputs were generated using custom written programs in R (R Core Team, 2012).

##### Hippocampal inputs during theta

To simulate the inputs to the CA1 pyramidal cell we focused on the population activity of the presynaptic CA3 pyramidal neurons and ignored inputs arriving from elsewhere, including the entorhinal cortex. A single CA1 pyramidal neuron receives *N*_syn_ ≈ 20000 excitatory synapses (Megias et al., 2001), and ≈ 10% of the presynaptic CA3 cells have place field in a typical environment (Alme et al., 2014; Hainmueller & Bartos, 2018). Therefore we simulated *N*_PC_ = 2000 place cells each exhibiting a single place field in the environment, while the activity of the other 18 000 cells was incorporated in the increased baseline firing rate of the simulated place cells.

In the simulations the animal was moving at a constant 20 cm/s speed on a 2 m-long circular track. Excitatory neurons had a single, idealised place field that showed phase precession relative to the ongoing theta oscillation (constant 8 Hz frequency). Phase precession was modeled using basis functions co-tuned to spatial location and theta phase. Specifically, we had *N*_basis_ = *N*_x_ · *N_ϕ_* = 160 Gaussian basis functions with *N*_x_ = 40 spatial and *N_ϕ_* = 4 temporal components uniformly tiling the space with standard deviation *σ*_x_ = 5 cm and *σ_ϕ_* = *π*/2 radians. The parameter w_*ϕ*_ was optimised numerically to match the phase precession data obtained from Skaggs et al. (1996) with the width of the place field being ≈ 30 cm (Figure S2I). To tile the space evenly, we shifted the parameter w_*ϕ*_ along the spatial dimension either randomly (Figure 1B-D) or evenly (Figure 1G-H, 50 cells with identical tuning, 40 ensembles). The average input rate of individual place cells along the track either chosen randomly from a gamma distribution with shape and rate parameters *α* = 6 and *β* = 6 (random, Mizuseki & Buzsáki 2013) or was identical 1Hz for all neurons (uniform) corresponding to a 5 Hz presynaptic firing rate when we take the low release probability of hippocampal synapses into account (*p*_rel_ ≈ 0.2, Holderith et al. 2012). To model spatially untuned activity of non-active place cells the parameter *λ*_0_ was set to 1 Hz leading to the average input rate 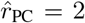 Hz for the simulated place cells. The total input rate of the excitatory inputs was thus 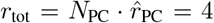 kHz matching the number of inputs a pyramidal neuron is expected to receive *in vivo*, 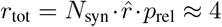 kHz, where 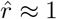 Hz is the average firing rate of a randomly chosen hippocampal pyramidal neuron (Mizuseki et al., 2012).

In both the uniform and the random condition we generated 10 different presynaptic population and simulated 16 trials with each population.

##### Hippocampal inputs during SPW

To simulate SPW events we embedded elevated hippocampal population activity of *T*_SPW_ = 100 ms duration in a low activity baseline state with independent presynaptic activity at a constant 0.8 Hz. During the SPW the inputs were driven by a simulated spatial trajectory corresponding to memory replay (Foster, 2017). The presynaptic cells had a spatial tuning and were also modulated by ongoing 150 Hz ripple oscillation (Buzsáki, 2015). Specifically, we used *N*_basis_ = *N*_x_ + *N_ϕ_* = 44 Gaussian basis functions with independent spatial (*N*_x_ = 40) and temporal (*N_ϕ_* = 4) components. The place fields had a diameter of ≈ 50 cm and were distributed uniformly across the entire track and the peak firing rate of each place cells were the same.

The average number of spikes fired by a rodent hippocampal pyramidal cell during an individual sharp wave event of *T*_SPW_ = 100 ms duration is *N*_sp_ ≈ 0.4 (Mizuseki & Buzsáki, 2013), leading to a *N*_syn_ · *N*_sp_ ·*p*_rel_ = 1600 synaptic events received by the postsynaptic cell. During the SPW a part of the previously experienced trajectory of the animal is replayed with an increased speed of *v* ≈ 6 m/s (Pfeiffer & Foster, 2013). In our 2m long environment ≈ 30% of the track is replayed in each SPW by the *N*_repl_ay ≈ 600 neurons having place fields overlapping with the replayed trajectory. We assume that these place cells are firing at an increased mean rate of *r*_replay_ ≈ 60 Hz and are thus responsible for *N*_replay_ · *r*_replay_ · *T*_SPW_ · *P*_rel_ = 720 synaptic events, while the remaining 880 inputs are uniformly distributed across the entire presynaptic population (19400 cells, out of which we simulated only 2000) by increasing the baseline firing rate of the place cells uniformly during the SPW period.

For the 240 inputs participating in synaptic clusters, the constant baseline firing was not increased, as these inputs represented input from a single presynaptic neuron. Instead, we selected the 240 of the 2000 not clustered synapses with location closest to the location of the clustered inputs, and these synapses received inputs with only baseline activity during SPW but no spatial tuning.

The firing rate of the inhibitory inputs targeting the perisomatic (dendritic) region switched from 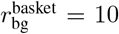 Hz 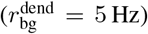 baseline rate to 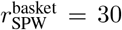 Hz 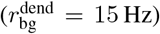 during SPWs, respectively. Both excitatory and inhibitory inputs were modulated by the 150 Hz ripple oscillation with the ratio between their peak / minimal firing rate being ~ 3 (Csicsvári et al., 1999). The depolarisation amplitude and the ripple modulation of the somatic membrane potential during SPWs in the biophysical model was consistent with data recorded from awake mice (English et al., 2014; Hulse et al., 2016).

We varied the starting position of the replayed trajectory in steps of 10 cm changing the overlap between the represented trajectory and the place field of the clustered neurons and analysed the average somatic response during SPWs in 16 independent trials.

##### L2/3 inputs

To generate *in vivo*-like synaptic inputs to a L2/3 pyramidal cell we simulated the activity of cortical and thalamic neurons in response to drifting grating stimuli, widely used to study neuronal coding in the visual system. Since the activity of these two populations is reasonably similar under these conditions (Durand et al., 2016), we did not treat them separately.

We modeled the response to visual neurons to gratings moving at 2 cycles/s with 16 different directions each shown for 1.5 s. The activity of each input was tuned to the direction and the phase of the stimulus with the distribution of firing rates, orientation selectivity, direction selectivity, response linearity (phase modulation depth divided by the mean firing rate; F1/F0 calculated as in Niell & Stryker 2008) and the width of the tuning curve matched to experimental data (Figure S5B; Durand et al. 2016; Niell & Stryker 2008). We used *N*_basis_ = *N_θ_* + *N_ϕ_* + *N_φ_* = 28 circular Gaussian basis functions with independent components responsible for the tuning to motion direction (*N_θ_* = 16), phase (*N_ϕ_* = 6) and gamma oscillation (*N_ψ_* = 6). Cells were divided into 16 groups based on their direction preference, and within each group we simulated 120 neurons with 5 different directional tuning patterns (including direction- and orientation-selective cells), 6 different preferred stimulus phase and 4 different phase tuning pattern, including simple cells (showing marked phase preference) and complex cells (having little phase preference). In the random condition (Figure S5D-E, random inputs) the orientation and stimulus phase preference of the cells were selected randomly with uniform probability and their average firing rate was chosen randomly from a gamma distribution with shape and rate parameters *α* = 2 · 4.17 and β = 2. In the uniform condition (Figure S5, except panel D-E, uniform inputs, F-J), each presynaptic input represented a unique combination of these features with identical mean firing rate 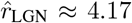 Hz. The total excitatory input rate of the L2/3 model neuron was around 8 kHz.

Inhibitory inputs had little orientation tuning, showed complex cell-like phase preference and had a mean firing rate of ≈ 30 Hz. We generated 10 different presynaptic populations with either uniform or random condition and simulated 16 trials with each population.

#### Decomposition of response variance

The subthreshold membrane potential response *r*(*x, k*) of the postsynaptic neuron at location *x*^*^ in trial *k* can be approximated by the sum of stimulus-dependent and dendritic factors, each factor potentially contributing both to the tuning curve and to trial-to-trial variability of the responses:

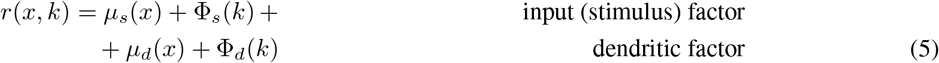

where *μ_i_*(*x*) captures the location dependent change in the mean response and Φ_*i*_(*k*) is the drive fluctuating from trial-to-trial that is associated with the given factor. Specifically, Φ_*s*_ is associated with fluctuation in the total input (presynaptic spike counts) arriving to the cell while Φ_*d*_ denote fluctuations in the input related to the location of the activated synapses within the dendritic tree brought about by passive (e.g. cable filtering) and active (e.g. location dependent activation of NMDA receptors) mechanisms as well as functional synaptic clustering (interactions between local active inputs).

We defined these factors to have a linear effect on the somatic response (membrane potential), so biophysically these factors correspond to currents flowing into the soma of the neuron. Note, that these factors do not correspond to separate biophysical processes: During the sustained synaptic background activity, the activation of a single additional synapse contributes to both dendritic and input factors as it increases the total input count (input factor) but the local depolarization it causes also modulates input current flowing though the neighbouring synapses (dendritic factor). Also note, that in general, these factors are not independent of each other, as larger input strength (Φ_*s*_) is potentially associated with larger NMDAR current or stronger interactions within the clusters (Φ_*d*_).

These factors can not be directly measured in real neurons or in even a biophysical model. Therefore, instead of pinpointing the effect of these factors on a trial by trial bases, we were aiming at identifying their average contribution to the neuronal response variability. To achieve this, we derived a novel analysis technique, the *decomposition of neuronal response variance* where we selectively modified the contribution of the individual factors and measure their effect on the response in a biophysical model.

To estimate the contribution of the specific factors, we performed 4 different type of simulations. In the first scenario we used random inputs and random synaptic arrangements and thus all factors were present and their overall contribution to the response variance was measured (R, random case).

In the second scenario, which we called the *uniform* input scenario *μ_s_*(*x*) = *μ_s_*, so the mean total input was independent of the position, and all fluctuations in the response were caused by either trial-to-trial variability or systematic changes in the dendritic factors. Moreover, we assume that the dendritic factors were identical in the uniform and in the random case, that is, we assumed that changing the input heterogeneity did not change the contribution of dendritic factors. This is a critical assumption that is not justified in most cases (e.g. when large, local fluctuation in the inputs can boost dendritic nonlinearities), but can be a reasonably good approximation when changes in the input strength are relatively small. Therefore we used this analysis only when the arrangement of the synapses was random or balanced (i.e., not clustered, see below), and therefore we did not expect large, systematic fluctuations in the synaptic drive on any given dendritic branch. In the uniform scenario Equation 5 reads as:

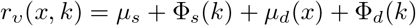

where the lower index in *r_ν_* indicates the condition (*ν* = ‘uniform’).

In the third simulation type, named as *fully* balanced we removed the dendritic factors by arranging synapses regularly throughout the entire dendritic tree. Instead of entirely eliminating the effect of dendritic processing, we only equalised its effect on the differently tuned inputs, so that dendritic components did not add to the variability of the neuronal tuning. This way we could selectively manipulate the effect of dendritic factors on the somatic response, without changing the contribution of the input factors. In the fully balanced scenario the systematic variability in the tuning was caused by input components, and we assumed that the contribution of the input components were identical in the fully balanced and the random case. In the fully balanced scenario Equation 5 simplifies to:

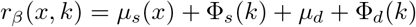

where the lower index in *r_β_* indicates the scenario (*β* = ‘balanced’).

As a control, we performed a fourth simulation type, where both the dendritic and input factors were eliminated. In this case we expected the postsynaptic tuning to be flat, as all variability was caused by trial-to-trial fluctuations:

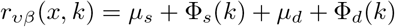

We denote the trial-to-trial variance of one component of the drive with 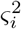, with *i* = {*β, ν*} coding for trial type, and assume that they do not change between the different scenarios. The correlation between the trial-to-trial components effect of the somatic and dendritic factors is denoted by *ρ_sd_*. The trial-to-trial variance of the response is expected to be nearly identical in all scenarios:

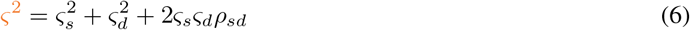

where the quantity in orange can be measured in a biophysical model. Confirming our assumption that changing the input and the synaptic arrangements does not change the trial-to-trial variability, we found that *ς*^2^ was similar in all four scenarios in the biophysical model of both the hippocampal (Figure 1J) and the visual cortical (Figure S5E) neuron.

When we average over *K* trials, the variance of the mean response (the estimated *tuning curve*) at a particular location will be *ς*^2^/*K*. Denoting the variance of the location dependent response by 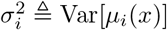, the variance of the estimated tuning curve in the four scenarios can be written as:

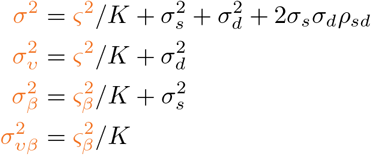

We estimated the effect of the individual factors by measuring tuning curve variances 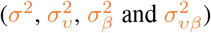 in the four different scenarios (Figure 1K).

#### Data analysis

To calculate the tuning curve from the recorded somatic membrane potentials we first detected action potentials (AP) as positive crossing of a threshold of *V*_th_ = −30 mV. We obtained the subthreshold response by replacing the APs by the voltage before the start of the AP, defined as the first point when the derivative of the voltage exceeded *η* = 5 mV/ms before the spike. The end of the AP was defined as the time when the membrane potential fell below threshold.

Next, we filtered the raw subthreshold response with a Gaussian kernel which removed the oscillatory component from the sVm. The width (SD) of the Gaussian kernel was *σ*_theta_ = 100 ms in the CA1 cell during theta, *σ*_ripple_ = 2.5 ms during SPWs and *σ*_gamma_ = 100 ms in the case of the L2/3 cell. We applied a similar Gaussian filtering with *σ*_theta_ = 100 ms to the input spike counts when we evaluated the input variability (Figure 1B,G,I and Figure S5D).

Finally the tuning curve was obtained by averaging the filtered subthreshold sVm responses across 16 trials with identical presynaptic firing rates but random synaptic events. The variability of the tuning curve was calculated as the variance of the points of the tuning curve along the track. The trial-to-trial variability was calculated by first computing the variance of the 16 trials and then taking the average across the track. The response integral during theta stimulation was calculated by first subtracting the baseline Vm defined as the mean sVm in the last 2000 ms from the individual filtered responses, and then integrating the baseline shifted response over time. The SPW amplitude (Figure 3B,E) was defined as the difference between the maximum of the average filtered somatic membrane potential (*V*_resp_) during the SPW and the minimum of *V*_resp_ before the SPW. The response amplitude (Figure 3G) was the difference between the SPW amplitude with maximal and minimal overlap between the replayed trajectory and the place fields of the clustered neurons.

### DATA AND SOFTWARE AVAILABILITY

The code used for simulating the biophysical model, generating the inputs and simulating and fitting hLN models can be downloaded from the online repository (https://bitbucket.org/bbu20/clustering).

## Supplemental Figures

**Figure S1.**
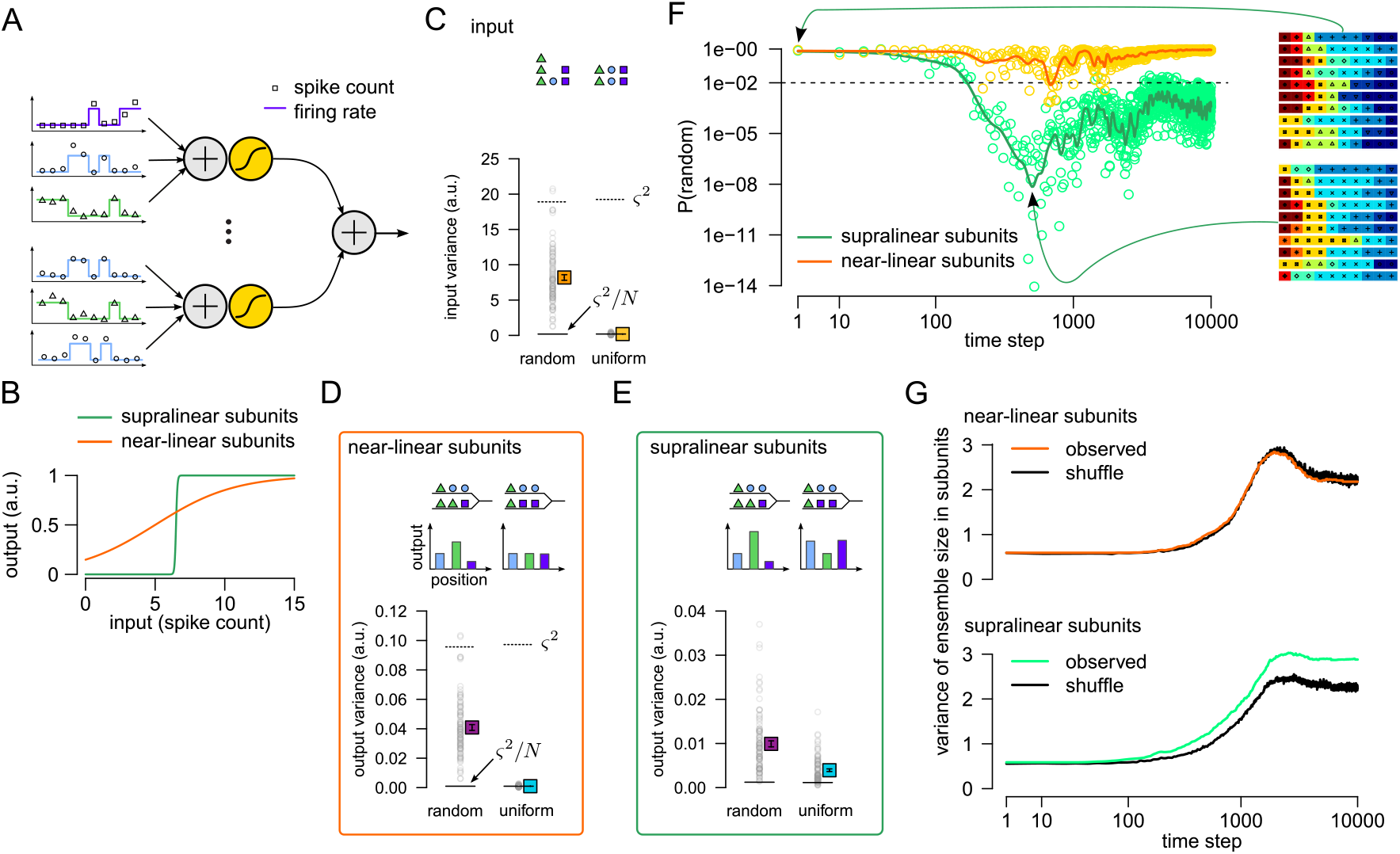
Global plasticity can lead to functional clustering. (A) Input integration in a simplified neuron model. Left: Presynaptic neurons (*N* = 100) were divided into non-overlapping ensembles with winner-take-all dynamics, representing groups of place cells with non-overlapping place fields. The firing rate (colored lines) of the ensembles switched between a low (1 Hz) and a high activity (10 Hz) state, and the activation of a particular ensemble corresponded to a specific spatial location. Spike counts (open symbols) were sampled from a Poisson distribution with the given firing rate in 100 ms time bins, corresponding to theta cycles. Right: Each of the 10 dendritic subunits (only 2 is shown for clarity) first integrated the incoming spike counts (+ sign) and then applied a static nonlinearity (yellow sigmoidal function) modelling dendritic spikes. The output of the neuron was the sum of the inputs from all branches. (B) By changing the parameters of the sigmoid subunit nonlinearity, *f*(*x*) = 1/(1 + exp(−*ζ*(*x* − *θ*))), we could interpolate between near-linear (orange) and supralinear (green) integration. Parameters were *θ* = 5, *ζ* = 0.35 for the near-linear and *θ* = 6.5, *ζ* = 20 for the supralinear subunits. (C) Variability in the mean total input spike counts across different locations (ensemble activation) when synapses are selected randomly from the 10 ensembles (left, random, see schematic on the top) and when the same number of inputs are selected from each ensemble (right, uniform). Horizontal lines indicate the expected variability of the mean input spike counts (*ς*^2^/*N*) and the trial-to-trial variability of the inputs (*ς*^2^). Grey circles represent *N* = 100 independent simulations, colored symbols indicate the mean ± SE. (D) Variance of the postsynaptic tuning curve (mean output across different locations) with near-linear subunits (orange line in panel B) reflects variability of the inputs. When there is no systematic variability in the input (see the insets showing two subunits with inputs from 3 ensembles) the tuning curve is flat, i.e., its variance is identical to the variance expected from trial-to-trial variability, *ς*^2^/*N*. (E) Same as D with nonlinear subunits (green line in panel B). The measured variability was larger than *ς*^2^/*N* even in the uniform case as subunit nonlinearities boosted clustered inputs (inset). (F) Simulation of structural plasticity in the model. The probability that the observed connectivity pattern was consistent with a random innervation in the case of linear (yellow) and nonlinear (green) subunits during the simulation of structural synaptic plasticity. The probability of random connectivity decreased markedly during the first 500 time steps, retaining a few dominant ensembles only in the case of nonlinear subunits (green), whereas the connectivity remains random when subunits were linear (yellow). Synaptic clusters emerged gradually from the initially random connectivity pattern as the output of the cell was biased towards ensembles organised into small synaptic clusters when subunits were nonlinear. During the global plasticity synapses not driving the cell were replaced by inputs from randomly selected ensembles that could stabilize if the same ensemble participated in at least one cluster. Inset: Connectivity patterns between presynaptic ensembles (color coded) and the dendritic subunits (horizontal rows) in one typical neuron at the beginning of the simulations (top) and at the most clustered stage (bottom). The degree of clustering decreases later when the number of different input ensembles decreases. (G) Variance of the cluster size was larger than in the shuffled control in the model with supraliner subunits, indicating the presence of functional clustering.

We simulated structural plasticity in the model by assuming that each synapse is associated with an abstract factor, *ϕ_i_*(*t*), representing biochemical processes stabilising the synapse. Initially *ϕ*(0) = 10 for all synapses. When *ϕ_i_*(*t*) ≤ 0 the synapse was replaced by an other input selected randomly from the 10 presynaptic ensembles. When the postsynaptic cell was active, *ϕ_i_* is increased by *α* = 5 for all active inputs and decreased for all inactive inputs by *β* = 1. We simulated plasticity in 25 independent cells fro 10 000 time steps. The P-value was calculated with a chi-square test where the random distribution of the largest cluster size was estimated by shuffling the connectivity patterns between the subunits of a given cell at each time step. Green and orange lines indicate moving average. Note the logarithmic *x* and *y* axis.

**Figure S2.**
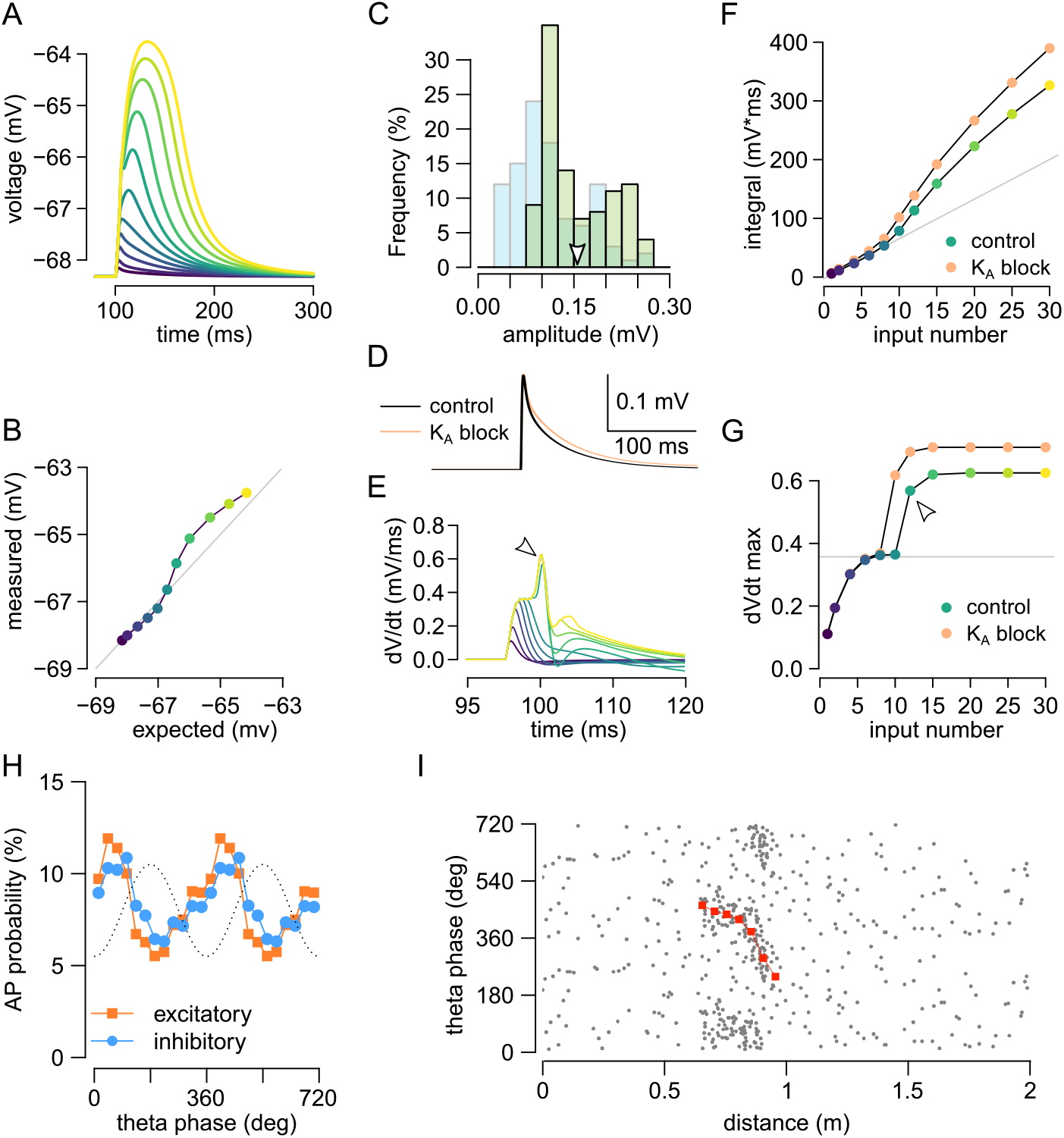
Dendritic integration and input statistics of the model CA1 neuron. (A) Somatic membrane potential in response to the activation 1-30 synapses on an oblique branch with *δ*t=0.3 ms. Color code is the same as in F-G. (B) Measured response amplitude as a function of linear expectations. Color code is the same as in F-G. (C) Histogram of the amplitude of the somatic response to the activation of 100 synapses distributed randomly along the apical trunk at 50-300 *μ*m distance from the soma (green, directly comparable to experimental data in Magee & Cook 2000) or along the entire dendritic tree (light blue). Arrowhead indicates the mean of the trunk synapse amplitudes. (D) Example of the somatic response to the activation of a single synapse in control conditions (black) and under partial blockade of dendritic A-type K^+^ channels. A-type potassium channels regulate the duration of EPSPs but not their amplitude. (E) Derivative of the somatic responses shown in panel A indicating the presence of local Na^+^ spikes for sufficiently strong stimuli (arrowhead pointing to the spikelets). Color code is the same as in F-G. (F) The integral of the somatic response as a function of the number of stimuli. Grey line shows linear expectation fitted to the first 5 datapoints. Orange dots show that the nonlinearity was larger when A-type K^+^ channels were blocked. The threshold for NMDAR nonlinearity was around 10 stimuli. (G) The peak amplitude of the derivative of the somatic response showed a step-like increase when local Na+ spikes were generated (arrowhead). The threshold for dendritic Na+ spikes was similar to the threshold for NMDAR nonlinearity (F). The amplitude of the somatic dV/dt was increased when A-type K^+^ channels were blocked (orange). (H) Theta modulation of inhibitory (blue) and excitatory inputs (orange) to the CA1 model neurons during the theta state. For similar experimental data see Fig 2J of Grienberger et al. (2017). (I) Phase precession of excitatory inputs. Grey dots indicate single spikes and red symbols show circular mean phase. Note that each spike is shown twice at *ϕ* and *ϕ* + 360°. Also note that the high background firing outside the place field accounts for the activity of ~10 cells not having place field on this track and thus not explicitely simulated. For similar experimental data see Fig 7 of Skaggs et al. (1996).

**Figure S3.**
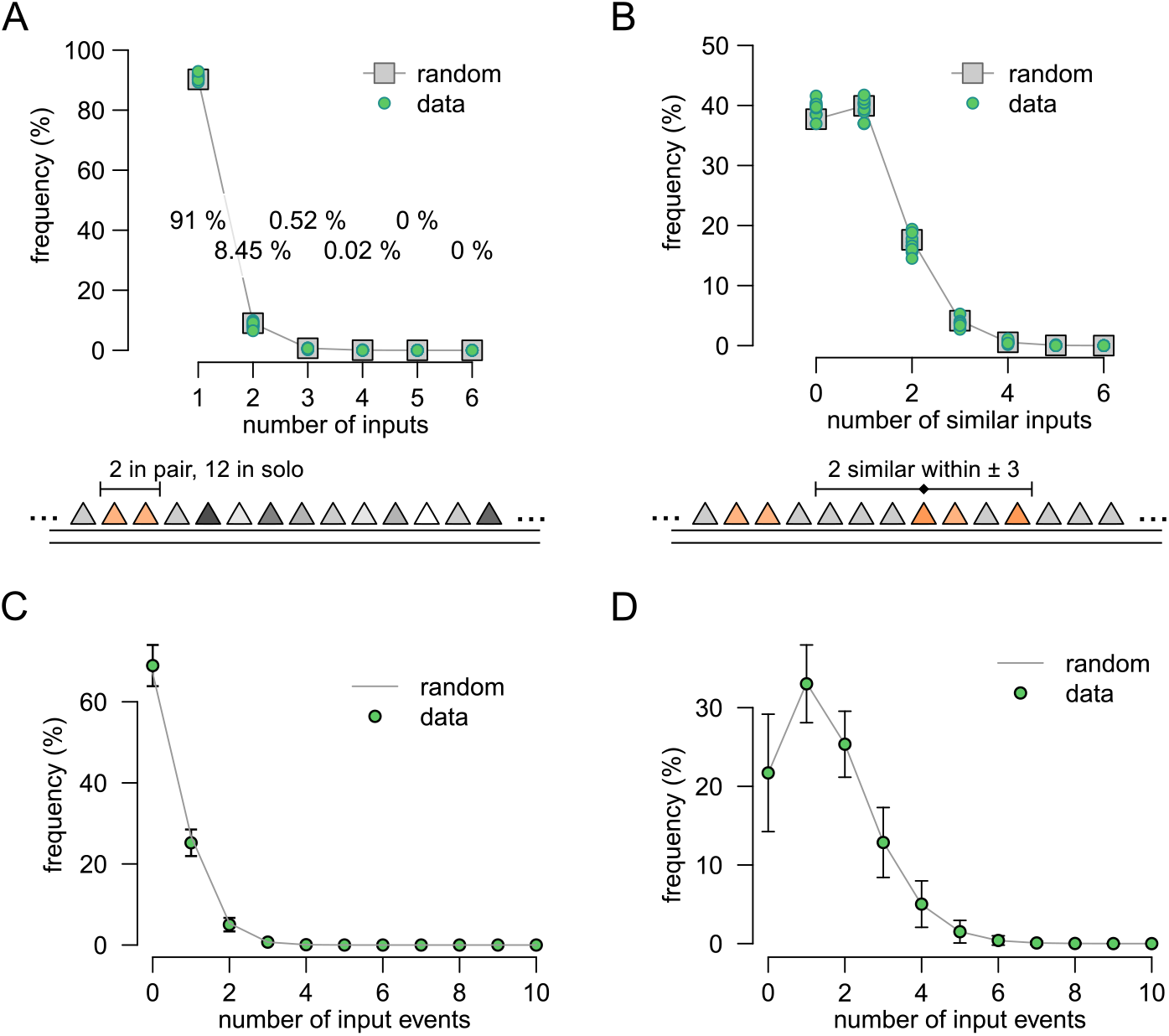
Synaptic clusters with unstructured innervation. (A) Distribution of the number of neighbouring inputs along the postsynaptic dendritic branch of the biophysical model (inset at the bottom) with identical place field location (green, n=10 different connectivity patterns). The data is consistent with a pattern expected from random innervation (grey) and indicates that ~ 8% of the inputs occurred in pairs but larger clusters contained less than 1% of the inputs. (B) Distribution of the number of inputs with similar tuning within the 6 neighbouring inputs (inset at the bottom). Here we used a different measure of functional clustering than in panel A: First, similarly tuned inputs could be separated with unrelated inputs. Second, we allowed a 15 cm distance between the peak of the presynaptic place fields. Even with this less conservative definition of functional synaptic clustering, clusters of 4 synapses were rare. (C) Distribution of the number of synaptic events arriving at 100 *μ*m long segment of a single dendritic branch within 10ms during theta state. With unstructured connectivity the typical number of input rarely exceeded 2 spikes/10 ms. (D) Same as panel C during SPWs, when the typical input could be as large as 4-5 spikes per 100 *μ*m dendrite within 10 ms.

**Figure S4.**
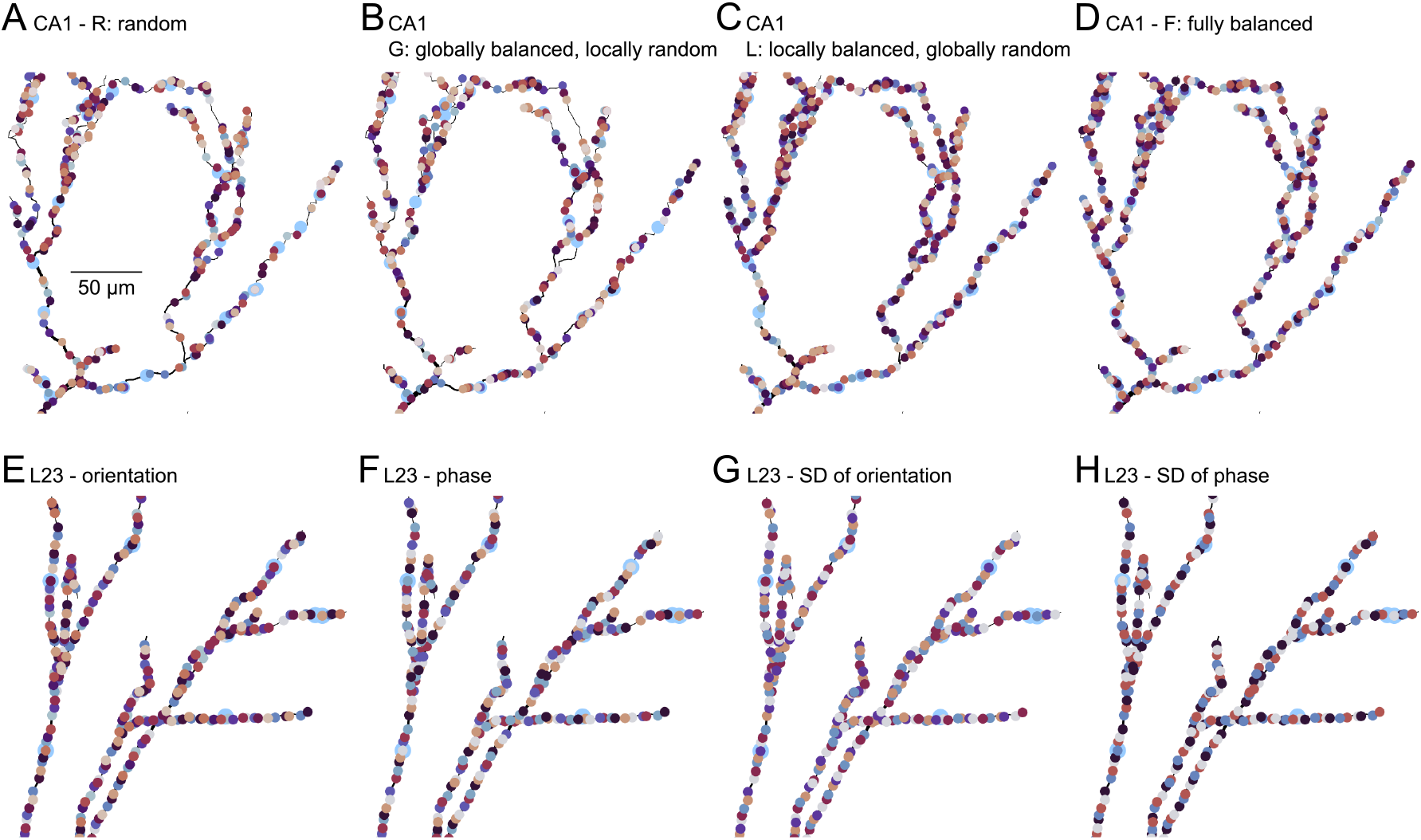
Synaptic arrangements in hippocampal CA1 (A-D) and visual cortical L2/3 cells (E-H). (A) Example for random synapse arrangement in a CA1 subtree. Colors indicate place field location, with distance in the color space matching distance between place field locations. Large blue circles indicate inhibitory inputs. (B) Example for locally random, globally balanced synapse arrangement. Synapses were first ordered globally into a single sequence mapped on the dendritic tree minimizing both small-scale functional clustering and large scale inhomogeneities. Next, synapses were permuted and their location were randomized within each branch, potentially generating small scale functional clustering but not large scale biases. (C) Example for globally random, locally balanced synapse arrangement. Synapses are first arranged randomly throughout the entire dendritic tree. Next, synapses within each branch are organised into a single sequence to remove small-scale functional clustering while retaining large scale biases. (D) Example for fully balanced synapse arrangement. Synapses were ordered globally into a single sequence mapped on the dendritic tree minimizing both small-scale functional clustering and large scale biases. (E-H) The fully balanced arrangement in visual cortical L2/3 cell. Inputs in the visual cortex were higher dimensional as they were tuned to both orientation and phase and had different tuning curve width (i.e., simple cells had a narrow phase tuning while complex cells were widely tuned). Inputs were organized in an approximately balanced way with respect to all four variables. Color code indicates the tuning of the cells with respect to the *N_ϕ_* = 16 different orientation preference (E), *N_ψ_* = 6 different phase preference (F), *N_ϕ_* = 5 different orientation tuning width (G) and *N*_Ψ_ = 4 different phase tuning width (H). As {*N_ϕ_, N_ψ_, N*_Φ_, *N*_Ψ_} were not co-prime numbers, we could not use independent co-prime reordering to achieve a coherent rearrangement of *N_ϕ_* × *N_ψ_* × *N*_Φ_ × *N*_Ψ_ = 1920 inputs. (E) We used co-prime reordering of the 16 orientation preference classes using *α* = 5. The same orientation-preference sequence was repeated 15 times (240 inputs) after which the base sequence was shifted by 3 for the next 15 repeats followed by further 15 repeats with shifts {6, 1, 4, 7, 2, 5}. (F) The sequence {1, 3, 5, 2, 4, 6} was repeated 320 times. (G) The sequence {1, 4, 2, 5, 3} was repeated 384 times. (H) The sequence {1, 3, 2, 4} was repeated 15 times and {3, 2, 4, 1} was repeated 15 times and then the whole sequence was repeated 16 times.

**Figure S5.**
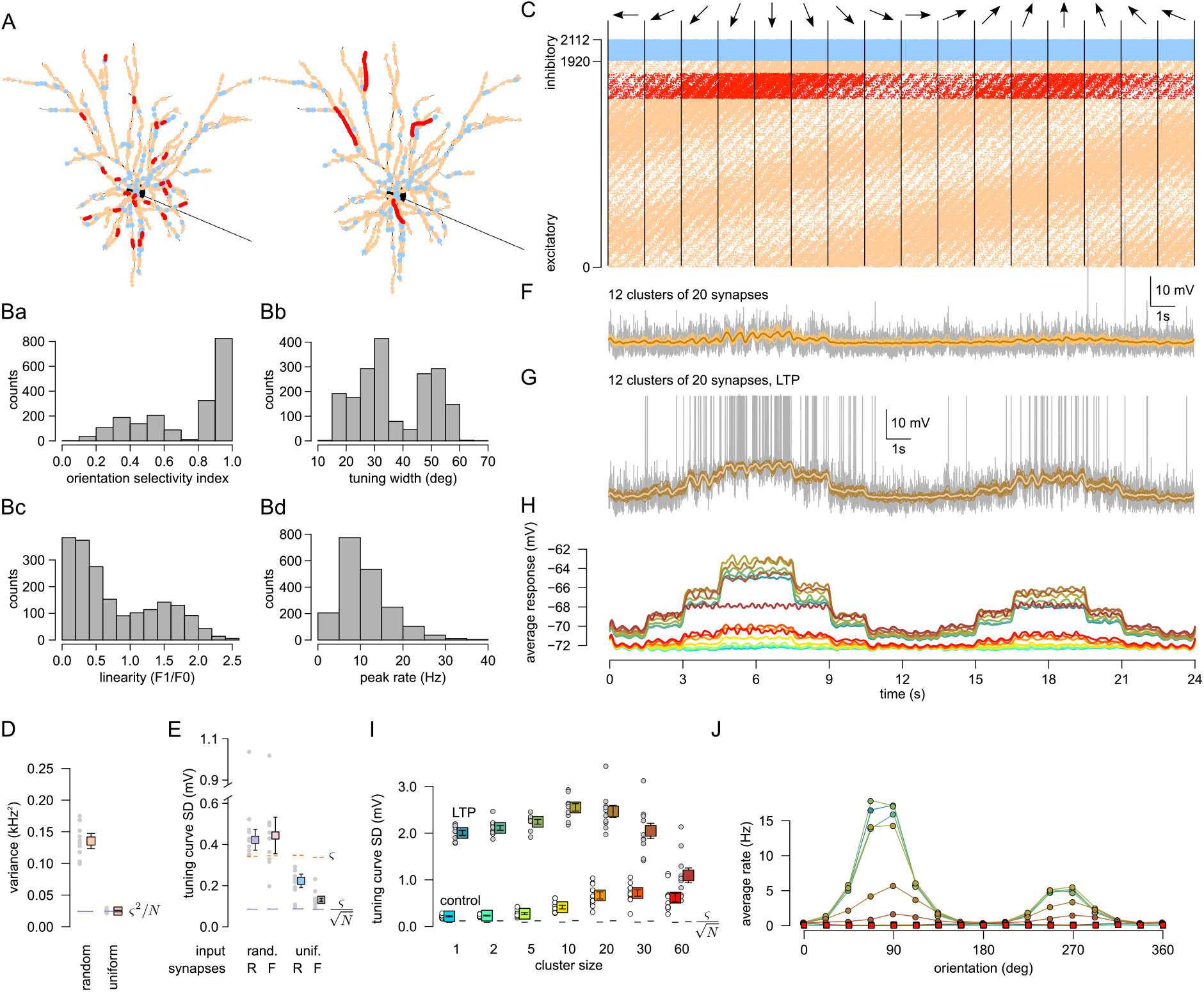
Effect of functional synaptic clusters in L2/3 neurons. (Related to Figure 1 and Figure 2.) (A) Placement 240 synapses (red) into 24 (left) or 4 (right) functional synaptic clusters together with the 1680 background excitatory (orange) and 192 inhibitory (blue) synapses on the model L2/3 neuron. (B) Distribution of orientation selectivity (Ba), tuning curve width (Bb), receptive field linearity (Bc) and the peak firing rate (Bd) of the inputs. Receptive field linearity was defined as the ratio between the amplitude of the oscillatory component of the response (F1) and the mean response (F0) to the preferred stimulus (Niell & Stryker, 2008). For comparable experimental data, see Fig. 4A, 4C 7A, and 8E of Niell & Stryker (2008), respectively. (C) Activity of excitatory (orange and red) and inhibitory (blue) inputs in response to simulated drifting gratings in 16 different orientations (arrows on the top), each shown for 1.5 s. Excitatory inputs were ordered according to their direction preference. A subset of the inputs responded to movements in the opposite directions (orientation selectivity) leading to the second, weaker band of spikes. Excitatory inputs preferring the down direction (red) were chosen for clustering. (D) Variability of the inputs. The variance of the mean filtered spike count is higher than (same as) the variance expected from trial-to-trial variability () when inputs are random (uniform), respectively. Solid lines indicate the variance expected based on trial-to-trial variability (*ς*^2^/*N*, purple). Symbols indicate mean±SEM. (E) Tuning curve standard deviation (SD) with random (R) and fully balanced (F) synaptic arrangements and with random or uniform inputs in the absence of systematic functional clustering. Purple line segments show the SD expected based on trial-to-trial variability 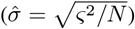 and orange dashed lines show trial-to-trial SD (*ς*). Since visual cortical inputs are tuned to multiple features, we could not entirely eliminate the effect of dendritic processing on the tuning curve even in the fully balanced configuration (tuning curve SD is larger than *ς*/*N* in the uniform, fully balanced case). Symbols in this panel show median and quartiles as data is highly non-Gaussian (note the outliers). (F) Example sVm response of the L2/3 neuron (grey), filtered responses for 16 trials (orange) and the tuning curve (brown) with 12 clusters of 20 synapses. Spikes were removed from the responses to obtain the filtered responses and the tuning curve. (G) Same as D with potentiated clustered synapses. (H) Average subthreshold response of the L2/3 postsynaptic cell showed an increased depolarization amplitude with clustering. Potentiation of the clustered synapses further increased the response amplitude. Color code is the same as in panel I. (I) Tuning curve SD as a function of synaptic clustering for control (bright colors) and LTP (dark colors) in 10 independent simulations (circles) and their means (color boxes) and SEM (error bars). Note that in E and I we show tuning curve SD instead of the variance (as in Figure 1), since the variance would further exaggerate the large differences between control and LTP. (J) Firing rate of the postsynaptic L2/3 cell as a function of orientation with different levels of clustering. Color code is the same as in panel I. The firing rate was near zero without LTP.

## Footnotes

* Here *x* can be a specific spatial location in the case of a CA1 neuron, but in general it is a point in a low dimensional manifold of the presynaptic firing rates.

